# Single-cell atlases of two lophotrochozoan larvae highlight their complex evolutionary histories

**DOI:** 10.1101/2023.01.04.522730

**Authors:** Laura Piovani, Daniel J. Leite, Luis Alfonso Yañez Guerra, Fraser Simpson, Jacob M. Musser, Irepan Salvador-Martínez, Ferdinand Marlétaz, Gáspár Jékely, Maximilian J. Telford

**Author notes:** Present address (Daniel J. Leite, Jacob M. Musser, Irepan Salvador-Martínez).

## Abstract

Pelagic larval stages are widespread across animals, yet it is unclear if larvae were present in the last common ancestor of animals or whether they evolved multiple times due to common selective pressures. Many marine larvae are at least superficially similar, they are small, swim through beating of ciliated bands and sense the environment with an apical organ structure. To understand these similarities, we have generated single cell atlases for marine larvae from two animal phyla and have compared their cell types. We found clear similarities among ciliary band cells and neurons of the apical organ in the two larvae pointing to possible homology of these structures suggesting a single origin of larvae within the clade analysed here (Lophotrochozoa). We also find several clade specific innovations in each larva, including distinct myocytes and shell gland cells in the oyster larva. Oyster shell gland cells express many novel genes which have made previous gene age estimates for trochophore larvae too young.

## Introduction

Indirect development via a small, pelagic, ciliated larva is found in at least 12 metazoan phyla (Fig. 1A). Ciliated larvae are generally understood to enhance dispersal in aquatic animals. The larval stage, which may last up to several weeks, is typically followed by a more or less rapid metamorphosis into a very different adult form. Six of the metazoan phyla with ciliated larvae - molluscs, annelids, nemerteans, brachiopods, phoronids and flatworms - are grouped within the super phylum of the Lophotrochozoa. Spiral cleavage is a second important shared characteristic among most of these phyla, supporting their monophyly within the Lophotrochozoa. The most iconic of these larvae, to which the others are often compared, is the trochophore larva found in the annelids and molluscs.

**Fig. 1.**
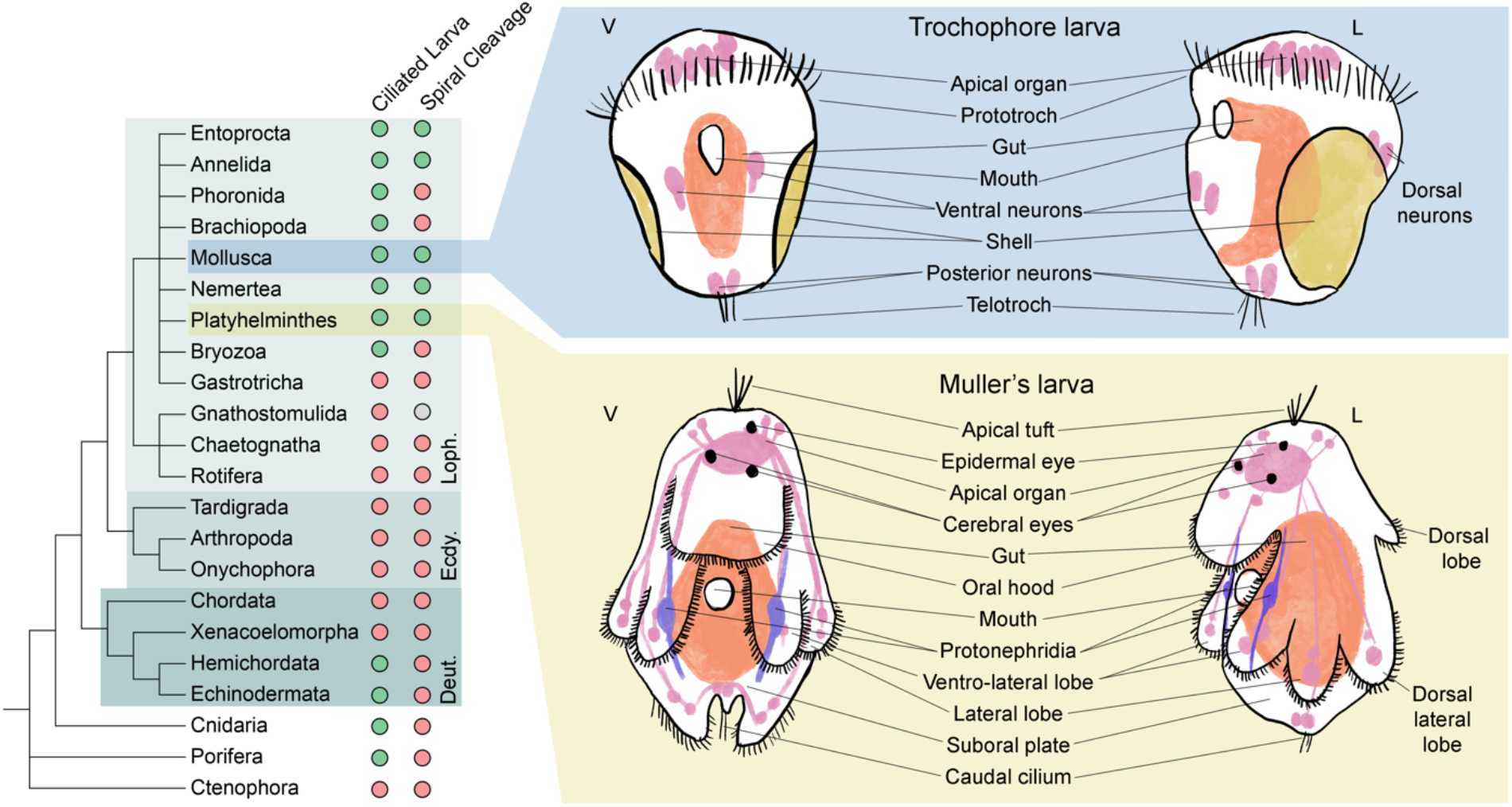
Larvae are a common feature of Metazoa. On the left, a phylogeny of animals shows widespread presence of ciliated larvae, especially among the superphylum Lophotrochozoa (Loph.). On the right, schematics of larvae presented in this study, top in blue trochophore larva of the Pacific oyster *C. gigas*, represented at both ventral (V) and lateral (L) view. *C. gigas* larvae at the trochophore stage lack an apical tuft of sensory cilia and paired protonephridia. In the bottom right in yellow, schematics of the Müller’s larva of the polyclad flatworm *P. crozieri* depicted in ventral (V) and lateral (L) view.

Trochophore larvae get their name from their preoral ciliary band or prototroch which is situated as an equatorial girdle dividing the animal approximately in two. The larval eyes are found on the anterior episphere and at the anterior pole of a typical molluscan or annelid trochophore there is a long tuft of sensory cilia called the apical tuft. Immediately below the prototroch is the mouth and additional ciliary bands may be found more posteriorly including a terminal telotroch (*1*). Internally, a canonical trochophore larva has an apical organ below the apical tuft, paired protonephridia and a larval gut (see Fig. 1B).

While lophotrochozoan larvae from phyla outside the molluscs and annelids have been considered to be derived forms of the canonical trochophore, these larvae have also been suggested to be cases of convergent evolution driven by similar selective pressures. The Müller’s larva of the polyclad flatworms shares with trochophores a locomotory ciliated band, anterior eyes, an apical tuft of cilia, an apical organ and protonephridia but in other ways differs from the trochophore (*2*). In particular, the body of the Müller’s larva is more complex; its ciliary band runs a convoluted path along the edges of eight lobes of the body (two ventro-lateral, two lateral, two laterodorsal, one oral and one dorsal) and is used for both filter feeding and for locomotion (see Fig. 1C).

The existence of both shared larval characters and clade-specific differences has prompted heated debates regarding lophotrochozoan larval evolution such as how often have larvae evolved, what are the adaptive advantages of larvae in different groups and what are the developmental and genetic underpinnings of the evolution of different larval types. Comparative efforts to understand the evolution of lophotrochozoan larvae have so far focussed on morphological, developmental and (sparse) molecular comparisons of putatively homologous shared structures (such as ciliary bands or apical organs) (*3*–*6*). For nearly all these larvae, however, we still lack a thorough molecular characterisation of larval anatomy and this would represent an essential contribution to this debate. We don’t yet know, for example, how many cell types they have, whether any of these are common between larval types and the extent to which different larvae may have unique cell types or larval organs.

Here, we have characterised the cellular component of two very different lophotrochozoan larvae: the trochophore larva of the pacific oyster and the Müller’s larva of the tiger flatworm *Prostheceraeus crozieri*. We have described multiple cell types and located these spatially in each larva using in-situ hybridization and hybridisation chain reaction (HCR), thereby revealing the complex anatomy of both larvae (*7*). By considering the phylogenetic ages of expressed genes in different cell types, we found that larvae appear to be made up of a combination of ancient and more recently evolved cell types co-existing in the larval body. Finally, we attempted to identify cell types that might be homologous between the trochophore and Müller’s larvae and have extended this comparison to the cell types of the primary larva of sea urchins (*8*). These comparisons highlighted molecular similarities between ciliary bands and apical organs of trochophores and Müller’s larva, which might indicate that these larvae are homologous.

## Results and Discussion

### Single-cell atlas of the pacific oyster trochophore larvae

We generated a cell atlas of the trochophore larva of the pacific oyster *e*.*g. -* a commercially important species readily available from fishmongers. This species has a chromosome-scale genome assembly and is becoming a popular model to study trochophore evolution (*9*–*11*). After in vitro fertilisation to produce trochophore larvae, we carried out four rounds of cell capture from three distinct dissociation experiments using the 10x Chromium scRNA-seq system. Initial shallow sequencing allowed us to select the best quality library (see Fig. S1) which was then sequenced further to obtain a final dataset of 8,597 cells expressing a total of 26,275 genes with a mean UMI of 30,000 per cell. Oyster larvae contain fewer than 500 cells, our data therefore represent approximately a 17-fold coverage of each larval cell suggesting that even rare cell types had been sampled. By applying dimensionality reduction, we identified a total of 37 distinct cell clusters including three ciliary cell clusters, two neuronal cell clusters, eight myocyte clusters, six shell related clusters, four hemocyte clusters and six proliferative cell clusters (as detailed below) (see Fig. 2A). To validate and to assign putative identities to the individual cell clusters, we selected specific gene markers for each and used a combination of literature searches, chromogenic *in situ* hybridisation (ISH) and hybridisation chain reaction (HCR) *in situ*s (*7*) (see Fig. 2B).

**Fig. 2.**
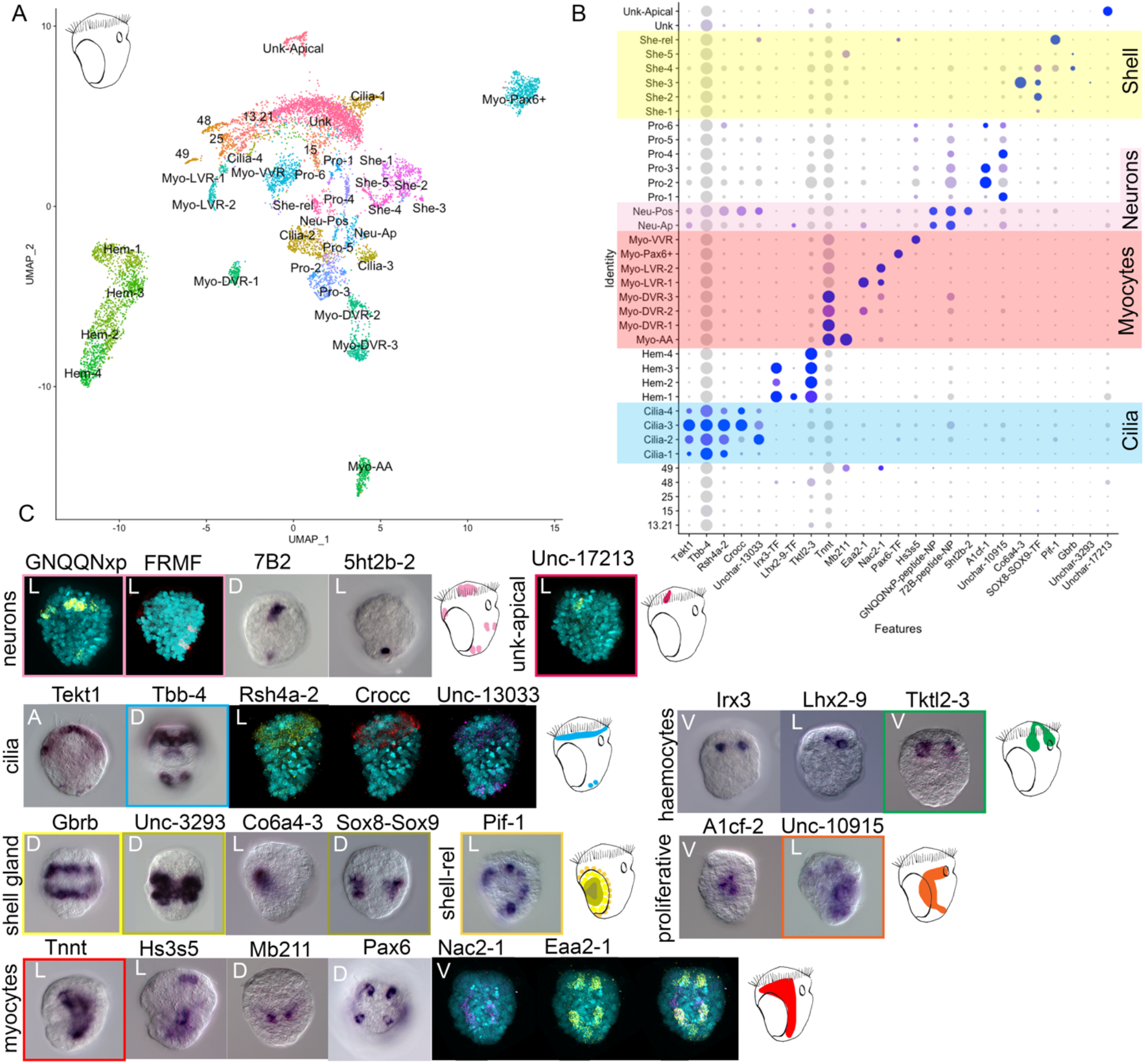
Cell atlas of the trochophore larva of *C. giga*s. A) UMAP showing cell clusters for the oyster larvae B) Marker gene expression for cell clusters, yellow highlights the shell gland clusters, pink the neuronal clusters, red the myocyte clusters and blue the ciliary clusters. C) ISH of the marker genes shown in B with schematic of expression in the larva. A: apical view, P: posterior view, V: ventral view, D: dorsal view, L: lateral view with mouth on the right.

#### Oyster trochophores have two ciliary bands, a reduced apical organ and no protonephridia

A striking feature of most marine invertebrate larvae is their ciliated bands, which have been proposed to be homologous across larval types (*3, 12, 13*). We recovered the expression of the ciliary markers *B-tubulin* and *tektin* (*14*) in four putative clusters of ciliary cells, which ISH and HCR stainings of selected marker genes showed were all localised in the prototroch (see Fig. 2C). In contrast, only markers of cluster Cilia-2 are expressed in both the telotroch and the prototroch (see Fig. 2B and 2C). The cluster Cilia-2 is characterised by the expression of several specific transcription factors (TFs) including *ASCL4, Pax2/5, Dbx, Gsc2* and *FoxL1*, while the TFs *PLSCR1-1* and the gene rootletin (*Crocc*) are specific to clusters Cilia-3 and Cilia-4. In general, the four ciliary clusters have 59 markers in common. The genes *lophotrochin* and *trochin*, recently suggested to be spiralian specific and shown to be expressed in the ciliary bands of various lophotrochozoan larvae (*13*), are expressed in clusters Cilia-1 and Cilia-3. Another eight of the spiralian-specific genes described by Wu and colleagues are also expressed in ciliary clusters (see Fig. S2). At the stage analysed here the larva does not have an apical ciliary tuft. To summarise, while we might have expected either that different ciliary bands were made of the same cell types or that prototroch and telotroch would have of a combination of four cell types we find that the small telotroch is made of a subset of cell types found in the prototroch.

A second feature typical of trochophore larvae is their apical organ. Previous experiments using immunocytochemistry against FMRFamide, serotonin and vesicular acetylcholine transporter (VAChT) in the oyster larva have identified a small apical organ, a pair of dorsal neurons, a pair of ventral neurons and a pair of posterior neurons (*15*). To identify neurons in our scRNA data, we searched for cells expressing neuropeptide precursors as well as a suite of additional neuronal markers (Fig. 2, Fig. S3, for a full list of NPs see table S1 and S2). We found two clusters of potential neurons that express the neuronal markers *cac1a-1, sy65-1* and *sy65-2* and the proneuropeptides *GNQQNxP, allatotropin* and *7B2* (see Fig. 2C and Fig. S3). ISH and HCR for marker genes from these two neuronal clusters revealed that one cluster corresponded to neurons in the apical organ and in the dorsal part of the larva (Neu-Ap) and the second corresponded to posterior neurons (Neu-Pos)(see Fig. 2B and 2C). Apical only proneuropeptides are *myomodulin* (which is restricted to only 1 or 2 cells of the apical organ) (Fig. S3), *CCWamide, corazonin* and *allatostatin*. The TFs *prd6, awh2, hbn, tbx5, prox, delta, sox2, zic1/2/5/4* are also specific to anterior neurons. *Awh, hbn, delta* and *zic* are also found in the apical organ of the sea urchin larva as well as in the nervous system of several other bilaterians (*16*). Posterior neurons are characterised by the expression of *TRH, MIP* and *AKH-GNRH, pedal peptide 1, CCAP-peptide* and *whitnin-PKYMDT-proctolin precursors*. The expression of the transcription factors *barH2, prop, pou4* and *evx* is also specific to the posterior cluster. ISH revealed only two posterior neurons (see staining of *5ht2b-2* Fig. 2), two dorsal neurons and a very small apical organ comprising approximately six neurons (see expression of GNQQNxP Fig. 2C). We identified two ventral neurons by HCR against the *FRMFamide* precursor mRNA, however, these ventral neurons are represented by only eight cells in our scRNAseq data and these do not form a separate cluster (Fig. 2 *FRMF* staining).

Overall, the oyster larva has an extremely reduced apical organ, made of only five to six cells with similar transcriptional signature. Few other neurons (two dorsal, two ventral and two posterior) are scattered around the body. It is possible that the low degree of neuronal complexity is due to the larva’s small body size (∼50µm) and their short larval life. None of the clusters analysed showed protonephridial staining and classical protonephridial marker genes (*e*.*g*., *pou3, sall, lhx1/5*) are not expressed in any cluster (*17*). Protonephridia were also not found in previous studies using immunohistochemistry against acetylated tubulin (*15*).

#### Oyster larvae have multiple distinct muscle types, immune and proliferative cell types

A search for muscle and myocyte-related genes revealed a previously undescribed complexity in the muscular system of the oyster larva. A total of 1,935 cells expressed multiple myocyte and muscle markers (including *tropomyosin, troponin-t (tnnt), twitchin* and several *myosin chain proteins*) as well as various transcription factors (including several Gata and Fox family genes) (Fig. 2, Fig. S4). The myocytes could be separated into eight distinct clusters and these can be associated with five morphologically distinct muscles within the larva based on ISH stainings: anterior adductor muscles (Myo-AA), dorsal velum retractor muscles (Myo-DVR), ventral velum retractor muscle clusters (Myo-VVR), larval retractor muscles (Myo-LVR) and an enigmatic group of myocytes expressing *pax6* (Myo-Pax6+).

M*b211*, marker of the Myo-AA cluster, is expressed on either side of the hinge of the forming shells, consistent with the location of the anterior adductor muscles; these are a bivalve innovation that control the opening and closing of the shell plates (*18*)(Fig. 2B and 2C). Cells of the Myo-AA cluster co-express both *paramyosin* and *calponin*, a combination that is typical of mollusc catch muscles, which are able to maintain passive tension for long periods of time with minimal energy requirements. Catch muscles are found in other invertebrates such as insects, crayfish, nematodes and brachiopods; in bivalves this muscle is used to keep their shell closed (*19*).

ISH for the marker gene *tnnt* revealed expression in two almost symmetrical patches on the anterior part of the trochophore corresponding to the location of the dorsal velum retractor muscles (DVR) (*20*). The cluster-specific marker *hs3s5* showed expression in the ventral side in the region of the ventral velum retractor muscles clusters (VVR). The two markers *eaa2-1* and *nac2-1* highlighted a cross-like pattern on the dorsal part of the larva consistent with the position of the larval retractor muscles (LVR) (Fig. 2B and 2C).

Finally, we identified a cluster (Myo-Pax6+) expressing several myocyte markers such as *troponin C, caldesmon-like, tubulin-2-b-chain, myosin essential light chain, actin*, and *muscle LIM protein*. Cells of this cluster also expressed several so-called “eye master regulators” such as *pax6, eya* and *six1/2* however no opsin expression was identified in this cluster. ISH of the cluster specific marker *pax6* showed expression in cells scattered symmetrically across the larva (Fig. 2B and 2C). This is not the first instance where *pax6, eya* and *six1/2*, as well as other so-called “eye master regulators” are found in non-photoreceptor type cells – these same TFs are found in the larval hydropore canal and coelomic pouches of sea urchins; as part of kidney development and the specification of somitic muscle in vertebrates; as well as in several other tissues in amphioxus (*21*).

The oyster immune system starts developing at the trochophore stage (*22*) and we identified four clusters expressing haemocyte related genes including *thymosin-beta, flotillin-2* and the TF *tal-1* (*23*–*25*). ISH for haemocyte cluster markers (*irx3, lhx2-1 and tktl2-3*; Fig. 2B and 2C) show expression in two patches on either side of the developing gut, which appear to be connected anteriorly. We also identified four clusters that express proliferative markers such as *mago-nashi 2* (*mgn2*), *sumo3, pcna* and *CBX1* that play a role in stem cell proliferation (*26*–*29*). ISH for markers of these clusters (*a1cf-1* and *unchar-10915*) showed expression in the region of the developing gut. This is in line with the previous observation that, at the trochophore stage, the gut is still developing (*20*).

In contrast to the very simple nervous system, the oyster larva presents a complex muscular and haematopoietic cell complement, with several myocyte types that appear to be specific to bivalve molluscs. We also found that, at the trochophore stage, the larval gut is still undergoing differentiation and cells present in this area have a proliferative-like signature.

#### Shell gland cell types express novel genes

A key character of the phylum Mollusca is the shell. A search for previously described shell gland markers such as *tyrosinase* (*tyro*), *mantle protein* (*mp*) and *nacrein* (*manl-9*) highlighted six clusters. Specific markers for these clusters show concentric rings of expression in or around the two dorsally located paired shell glands (see Fig. 1B and 2C). Specifically, a marker for the clusters She-1 and She-2 (*Unchar-1938*) stains the inner area of the shell gland, the marker for cluster She-3, *co6a4-3*, stains the hinge between the two presumptive shells and a marker for She-4 and She-5 (*gbrb*) stains the outer layer of the shell gland. Finally, the marker of the cluster She-rel, *pif-1*, stains the outermost layer. It is possible that some of the inner cells are responsible for secreting the prodissoconch I (the two D shaped larval shells) and the more external cells will secrete the prodissoconch II (the veliger shell) at a later stage (*30*).

Our results show the existence of cell types involved in making shells that may represent relatively recent mollusc-specific innovations. Stage-specific bulk RNA-seq has recently been used to explore the expression of phylogenetically older and younger genes throughout the development of lophotrochozoan larvae. This analysis revealed a peak of expression of young genes in the trochophore stage of the pacific oyster as well as the scallop *P. yessoensis* (*10, 11, 31*). These findings were interpreted as indicating that the mollusc trochophore was an evolutionarily recent innovation and arguing against the idea that the trochophore larva is the phylotypic stage of molluscs (*10*). We extended this phylostratigraphic approach to a cell-type resolution by computing the phylogenetic ages of genes expressed in each characterised cell type of the oyster larva. We found that the expression of very young genes was restricted to the shell gland cell types, which therefore have the highest (youngest) transcriptomic age index (see Fig. 3A and 3B). Our result indicates that the peak in TAI observed in previous studies is likely an artefact caused by the innovation of a shell added to an older larval body. Alternatively, the shell producing cells may have undergone more rapid evolution than the rest of the larval body, producing a subset of cells with a TAI that is younger than the rest of the larva. Our result highlights the benefit of looking at gene expression at the cell-type level rather than in bulk. The lowest (oldest) TAI is found in cell types such as proliferative cells and haemocytes (Fig. 3A).

**Fig. 3.**
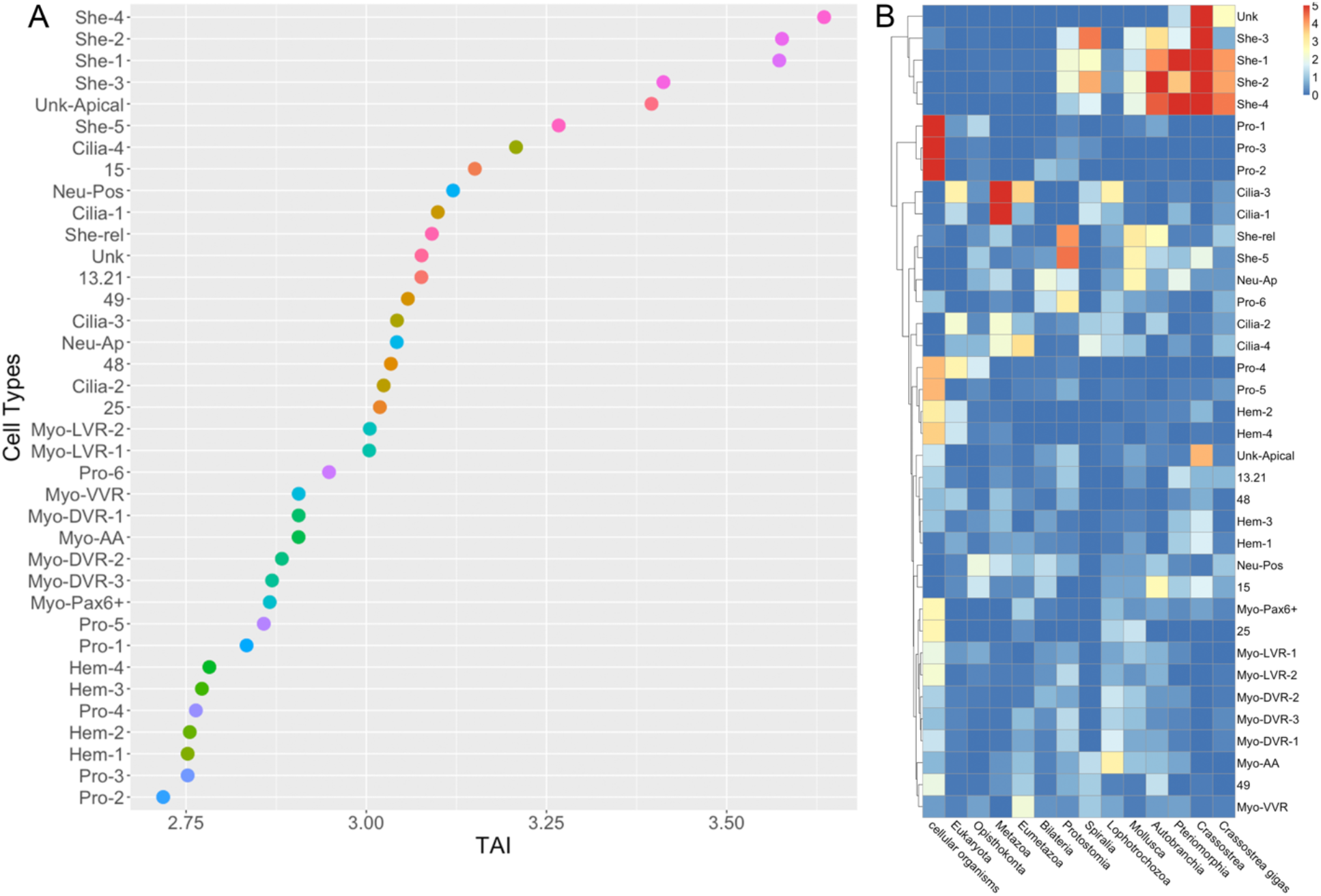
Gene age analyses in different cell types of the oyster larva show shell gland cells have a younger gene signature. A) Transcription age index (TAI) for different cell types, smaller TAI values correspond to “older” gene age. Gene age is inferred using a phylostratigraphy approach, then transcriptomic age index is calculated on the log transformed gene average expression per cluster. B) Phylostrata enrichment analyses per cell type. Phylostrata enrichment was computed using a hypergeometric test applied to the number of marker genes in each cluster per phylostrata compared to the global set of expressed genes.

### Single cell atlas of the polyclad flatworm Müller’s larva

To generate the Müller’s larva cell atlas, we carried out four rounds of cell capture from two separate dissociation experiments using 10x Chromium scRNA-seq. Initial shallow sequencing allowed us to select the best quality libraries which were then sequenced further to obtain a final dataset of 17,605 cells expressing a total of 33,305 genes with over 40,000 mean UMIs per cell (see Fig. s5). Müller’s larvae contain ∼2000 cells so our datasets represent an average of eight-fold coverage of each individual larval cell. Dimensionality reduction allowed us to identify a total of 51 distinct cell clusters; these include three ciliary clusters, eight neuronal clusters, two myocyte clusters, three neoblast clusters, nine gut related clusters, two clusters of rhabdites, five mesenchymal clusters, two cathepsin clusters and one cluster of protonephridia (Fig. 4A). To validate and assign putative identities to the individual cell clusters, we selected specific gene markers for each and used a combination of literature searches, HCR *in situs* and comparisons with the published cell atlas of the adult planarian worm *S. mediterranea* (*32*).

**Fig. 4.**
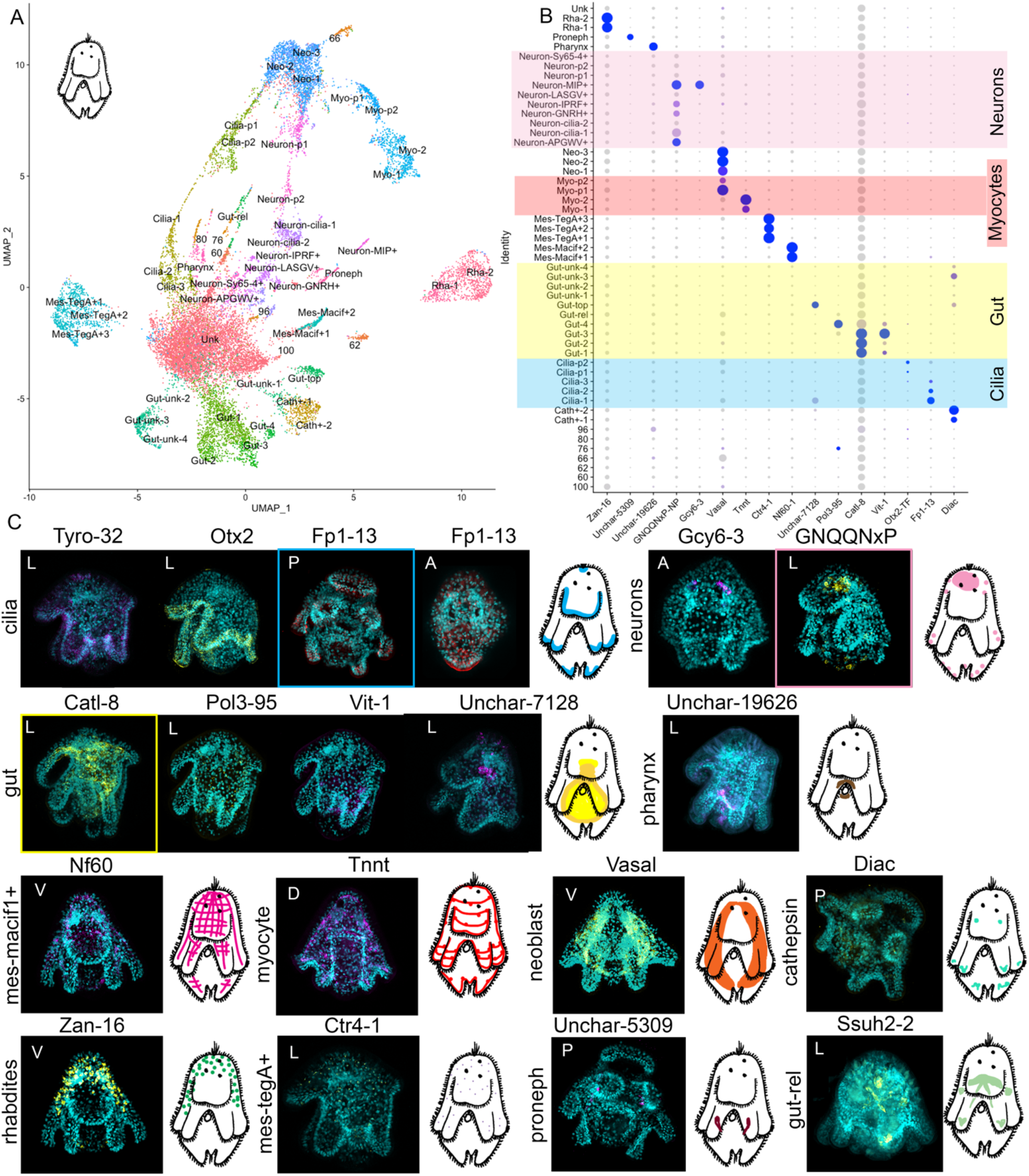
Single-cell atlas of the Müller’s larva of *P. crozieri*. A) UMAP showing cell clusters for the polyclad flatworm larvae. B) Marker gene expression for cell clusters, yellow highlights the gut clusters, pink the neuronal clusters, red the myocyte clusters and blue the ciliary clusters. C) ISH of the marker genes shown in B) with schematic of expression in the larva (highlighted gene/s used for schematic). A: apical view, P: posterior view, V: ventral view, D: dorsal view, L: lateral view with mouth on the left.

#### The Müller’s larva has simple set of muscle types and a complex gut

A search for myocyte markers, such as *troponin-i* (*tnni*), *troponin-t* (*tnnt*), *tropomyosin* (*tpm2*), *titin* and *paramyosin* revealed a total of four muscle clusters, two of which represent myocyte precursors (these cells also express neoblast markers such as Vasa). All the markers of Myo-1 (∼70 genes) are a subset of markers found also within the Myo-2 cluster. Myo-2 has an additional ∼350 specific marker genes. It is possible that these represent different myocyte cell-states and/or one of the two clusters is still undergoing differentiation. It is also evident on the UMAP that these two clusters are not separated from one another. The expression of the general marker troponin-c stains the larval muscles and resembles what has previously been shown using immunohistochemistry (2). The Müller’s larva thus has a considerably simpler muscle cell-type complement than the mollusc trochophore.

Contrary to what we found in the oyster, at least eight distinct clusters of cells had markers with expression in or around the gut. These include a cluster of cells near the top of the gut, highlighted by the (flatworm specific) cluster marker *Unchar-7128*, a cluster of cells localised in the posterior part of the gut, shown by the marker vit-1 and scattered cells in the gut lining (*pol3-95*). We also found a cluster (Gut-rel) containing cells located in an inverted “V” around the pharynx, above the dorsal part of the gut and in the ventro-lateral lobes (see expression of *ssuh2-2* Fig. 4). Finally, we identified a cluster of pharyngeal cells highlighted by the flatworm-specific marker gene *Unchar-19626*. Unlike the oyster trochophore larva, the Müller’s larva has a clearly differentiated gut made up of several cell types with specific expression signatures, and presumably distinct functions, similar to the planktonic larva of the sea urchin (8).

#### Trochophore-like features found in the Müller’s larva

The possible homology of the ciliary bands of flatworm larvae and those of trochophores has long been debated and their development in the two larval types shows both similarities and differences (*2, 33*). We found five clusters of ciliary band cells in the Müllers larva, two of these may represent differentiating ciliary precursors and the other three differentiated ciliary clusters. The three differentiated ciliary clusters share only 16 marker genes. Cluster Cilia-1 is characterised by the expression of hundreds of specific markers (for a full list see supplementary material), Cilia-2 is characterised by two specific caveolins (*cav1-8* and *cav1-9*) and the gene *entk* while Cilia-3 shares most markers with the cluster Cilia-1. HCR against the general markers *tyro-32, otx-2* and *fp1-13* show expression in the ciliated cells of the larval lobes as well as in the apical tuft (see Fig. 4). Similar to the myocytes, some of these clusters may represent differentiating cells rather than separate cell types.

Another typical feature of trochophore larvae, which we didn’t find in the oyster, are their excretory organs, the protonephridia. Protonephridial markers such as *pou2/3, hunchback* and *six1/2-2* allowed us to identify a cluster of protonephridial cells in the Müller’s larva (Fig. 3)(*17*). Cells from this cluster also express *cubulin* (specifically expressed in tubule cells of planarian protonephridia), *nephrin* (*nphn-1*) and *zo2* (*34*).

Apical organs have been proposed to be homologous between ciliated larvae (*4, 35*), for this reason we were interested in identifying them in our datasets and comparing their genetic signatures. In the Müller’s larva, HCR against a general marker for neurons (*GNQQNxP)* showed a large apical organ that sits around the ciliary eyes, and several scattered neurons elsewhere in the larval body (Fig. S3, 4C and 5). A combination of neuropeptides (NPs) and other neuronal markers allowed us to distinguish the presence of eight distinct neuronal clusters, each with its own signature combination of specific NPs (Fig. S3, for a full list of NPs see table S1 and S3).

We found a small cluster of neurons characterised by the expression of *IPRFamide* and *corazonin precursor genes. IPRFamide* precursor is expressed in two cells located apically as well as in one cell located in the suboral plate (Fig. 5 *IPRFamide* staining). A second cluster of anterior cells expresses *FRMFamide* and *LASGVamide precursor* genes as well as the TFs *Tbr1* (Fig. 5 *LASGVamide* staining). A third cluster contains *APGWamide* positive cells (Fig. 5 *APGWamide* staining) that are localised in large symmetrical patches laterally on the top part of the larva. Cells from this cluster are characterised by the expression of several other proneuropeptides: *APGWamide, GNQQNxP, calcitonin, allatostatin-A/buccalin, orexin/allatotropin* and *7B2* (Fig. S3). A fourth neuronal cluster contains *GNRH+* cells which show expression in a small patch near the apical tuft as well as in a couple of posterior neurons. Cells from this cluster also express the proneuropeptides *FVRIamide, FRMFamide, 7B2, XHFamide, NKY1* and *bursicon* (Fig. S3).

**Fig. 5.**
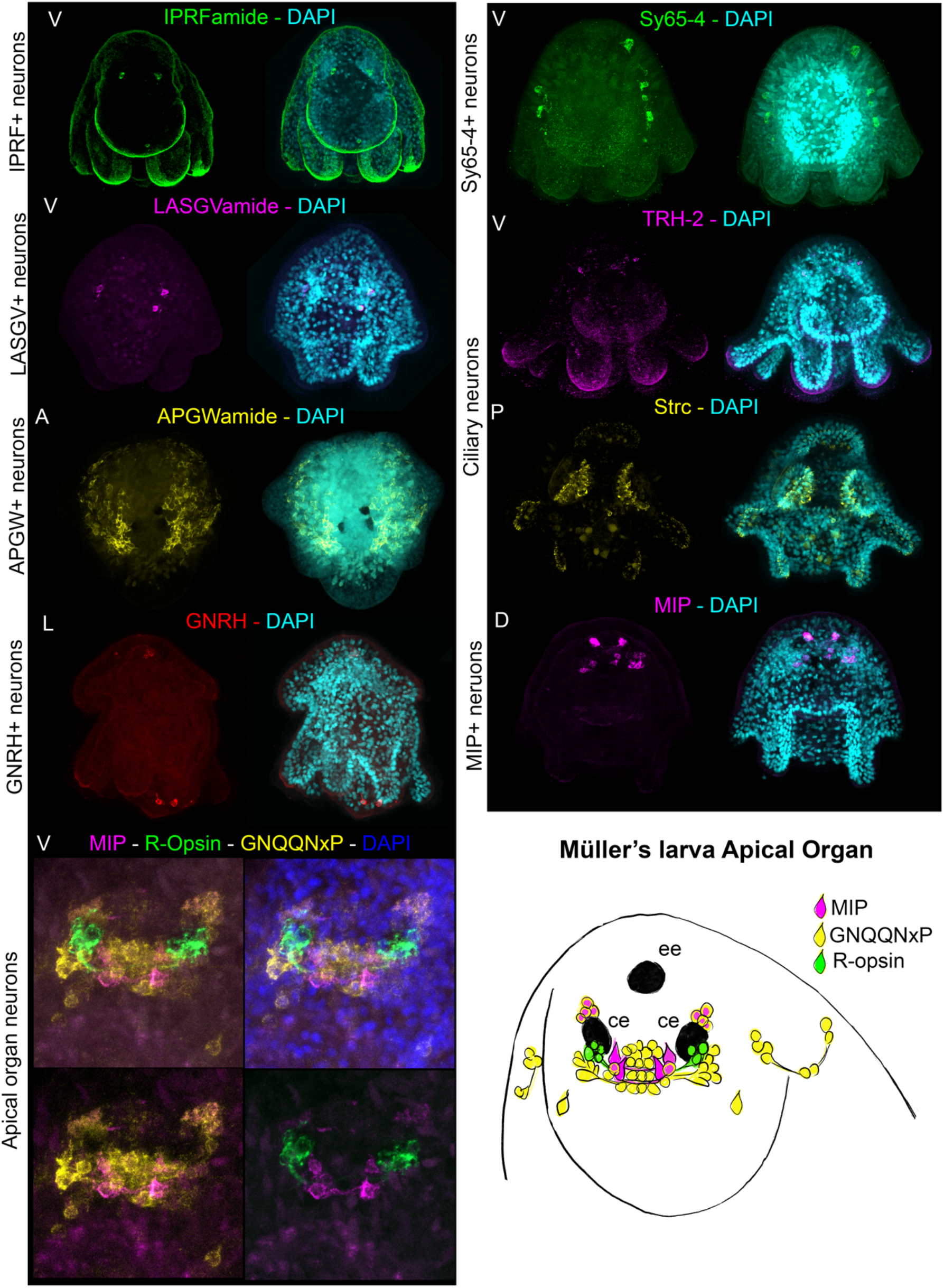
The complexity of the larval nervous system of the polyclad flatworm larva. HCR stainings of cluster markers and neuropeptides and schematic drawing of the Müller’s larva apical organ. All HCR stainings are maximum projections. A: apical view, P: posterior view, V: ventral view, D: dorsal view, L: lateral view with mouth on the left.

We also identified a fifth cluster of neurons that express several neuronal genes (including synaptotagmin (*sy65-4*), voltage-dependent calcium channel subunit alpha-1 (*cac1a*), voltage-gated sodium channel (*scna*) and a vesicular glutamate transporter (*vglu2*)) but no proneuropeptides (Fig. S3). Cells from sy65-4+ cluster also express the TFs *ETV4, repo* and *prox-1*. The HCR staining for the marker gene *sy65-4* highlighted several cells bulging into the epidermal layer on each side of the apical part of the larva (Fig. 5).

Specifically in the apical organ, we found a cluster of neurons expressing a cocktail of proneuropeptides (*MIP, GNQQNxP, LSDWNamide, 7B2*), several neuronal markers such as *synaptotagmin*, the acetylcholine synthesising enzyme *ChAT* and the *vesicular glutamate transporter* (v*glu2*) and the TFs *sox14, HLF-1* and *HLF-2* (Fig. S3). *HLF* is found in the apical organ of the sea urchin larva and in the nervous system of several other animals (*16*). The cluster specific marker *MIP* showed expression in two groups of cells posterior to the cerebral eyes, and in a cluster of cells anterior to these. These four anterior cells, connected in pairs by neuronal processes (see Fig. 5) are reminiscent of flask cells seen in the apical organs of other larvae (*36*). Moreover, the expression pattern of *MIP* resembles that observed in the annelids *Platynereis* and *Capitella* (*37*).

Two further clusters are of particular interest with regards to the potential homology of larval structures. These are identified as ciliary neurons and express several neuronal markers including *cac1a3, chat1, sy65-2* and *vacht-1* as well as the proneuropeptides *LSDWNamide* and *TRH2* (Fig. S3). These are likely to be the cilio-motor neurons described by Lacalli in another polyclad larva (*Pseudoceros canadensis*) and are also found in *Platynereis* larvae, where they are thought to be a larval specific character (*38*–*40*). Markers for these show expression in multiple cells in the ciliated lobes of the Müller’s larva (see Fig. 5) very similar to what has been seen in *Platynereis*. These markers are also expressed in a pair of apical cells. Equivalent cells are not found in the mollusc trochophore.

Our results show that the Müller’s larva of polyclad, described by some as a fairly derived larva, presents many trochophore-like characters such as ciliary bands, protonephridia, cilio-motor neurons and a complex apical organ. In the apical organ we find a specific subset of *MIP+* cells similar to cells found in other lophotrochozoan larvae (*37*).

#### Flatworm specific cell types in the Müller’s larva

We found several clusters of cells that appear to be flatworm specific including two clusters of secretory gland cells or rhabdites (identified with the marker *zan-16)*, cathepsin cells (identified with the marker *diac*), a cluster of cells that create a net-like structure in the mesenchyme (identified by the marker *Nf60*) and another cluster of scattered mesenchymal cells (*ctr4-1*). Some of these mesenchymal cells may act together as part of an adhesive organ (involving secretion) like the one found in the flatworm *Macrostomum lignano* with which they share the expression of the marker *macif1* (*41*).

We also found a large cluster of cells that expresses neoblast markers such as *vasa* and *piwi*. ISH for the gene *vasa* shows expression in the lateral mesenchyme of the animal. To clarify whether these were likely to be neoblasts and to confirm the identity of other clusters, we mapped our polyclad scRNA data to those of the planarian flatworm *Schmidtea mediterranea* using an unbiased approach: the self-assembling manifold mapping approach (SAMap) (*32, 42*). SAMap uses a measure of the similarity of expression of homologous genes to align single cell datasets across species (*42*). We found a good match between our hypothesised neoblast cells and the known planarian neoblasts as well as good matches between muscle cells, several gut clusters, cathepsin cells, protonephridia, and several neuronal clusters (Fig. S6). We also found that ciliary band clusters of the Müller’s larva matched the epidermis, protonephridia, and pharynx clusters of the adult planarian worms (all of which have cilia). The cells of the pharynx of the adult planarian did not match with those of the larval polyclad.

When looking at the transcriptomic age index of the polyclad flatworm larva, we found that some cell types, including two gut clusters, neuronal cells and some ciliary cells (Cilia-2 and Cilia-3) have the highest TAI values (i.e., express younger genes) and that neoblast and muscle precursors have lower TAI values (Fig. S7A and S7B). The ciliary cluster Cilia-2 on the other hand shows enrichment for Spiralian specific genes (Fig. S7B). While TAIs are not directly comparable between species, we did not find that putatively novel cell types of the Müller’s larva (cells unique to flatworms) express very young genes in the same way the oyster shell gland cells do.

### Which cell types are homologous in larvae of oysters, flatworms and sea urchins?

We have shown that the trochophore larva of the pacific oyster and the Müller’s larva of polyclad flatworms share several broadly similar cell types such as myocytes, proliferative cells, ciliated cells and neurons. To assess the potential homology of these cells in the two larvae, we used SAMap to compare all cells and all genes. We found that cell types such as myocytes, proliferative cells, ciliary cells and neurons broadly aligned across species (see Fig. 6A). We also found unexpected matches between the haemocytes of the oyster and the cathepsin cells of the flatworm and between the shell gland cells of the oyster and the Macif1+ mesenchymal cells of the flatworm. In planarian flatworms, *cathepsin+*cells include glial cells, pigmented cells and some cells with roles in phagocytosis (*43*); it is possible that flatworm cells with this latter function map with the oyster hemocytes. The match between oyster shell gland cells and flatworm-mesenchymal cells may reflect a common secretory role.

**Fig. 6.**
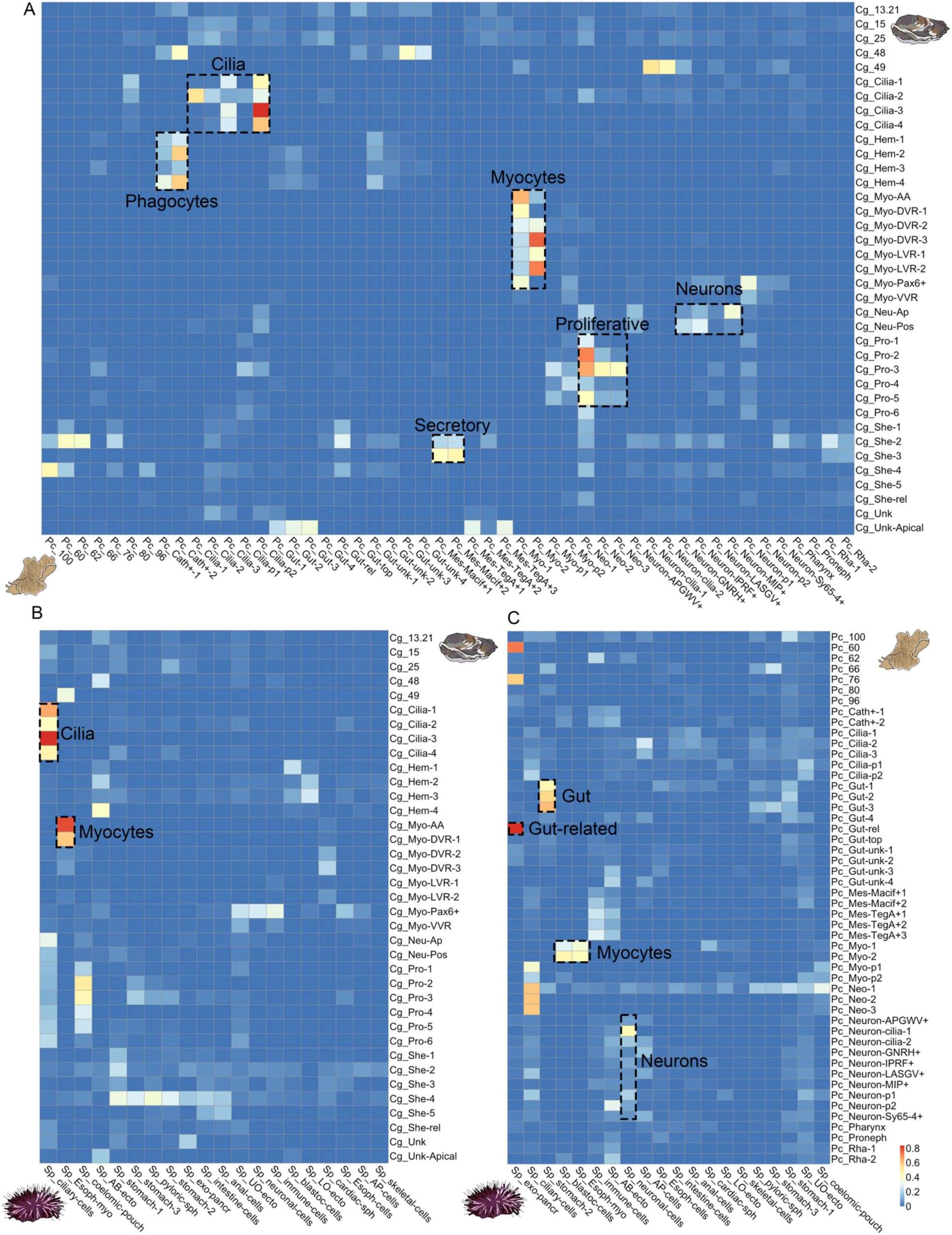
SAMap cell types alignment scores between invertebrate larvae. A) mapping of cell types between the trochophore larva of the pacific oyster (Cg) and the Müller larva of polyclad flatworm (Pc) B) mapping of cell types between the trochophore larva of the pacific oyster (Cg) and the pluteus larva of sea urchin (Sp) C) mapping of cell types between the Müller larva of polyclad flatworm (Pc) and the pluteus larva of sea urchin (Sp). Alignment scores are defined as the average number of mutual nearest cross-species neighbors of each cell relative to the maximum possible number of neighbors (*42*).

As ciliary bands and apical organs are both larval-specific characters, we explored the matches between the ciliary cells and between neurons of the two larvae by looking at co-expressed genes highlighted by SAMap. We found 133 oyster and 112 flatworm genes co-expressed between ciliary clusters (the difference in numbers is due to SAMap allowing one to many orthologs/paralogs matches). Many of these are proteins known to be involved in the ciliary apparatus such as *annexins, calmodulins, rootletins, centrins, calcineurins*, cilia and flagella associated proteins, *cubulins*, various *dyneins, enkurin, stomatin, tektins*, as well as several genes that remain uncharacterised (for a full list see Fig. S8). Only two TFs were co-expressed (CEBPB 2 and 3 and CEBP-G) however these have a broader expression and are not specific to ciliary bands (Fig. S8).

We surveyed the expression of all homologous TFs between the two species to see whether SAMap had missed genes due to their, usually low, level of expression (see Fig. S9). We found that several TFs not identified by SAMap (*Dbx, Gsc, Otx, Pbx, Zfhx* and *Pax6*; Fig. S9) are expressed in ciliary band cells in both flatworm and oyster. *Otx* is also expressed in the ciliary bands of larvae of the mollusc *Patella vulgata* (*44*), the annelid worm *Platynereis dumerilii* (*45*), the hemichordate *Ptychodera flava* (*46*) and in echinoderm larvae (*47*). We also found the lophotrochozoan specific gene *lophotrochin* in the ciliary cells of both the oyster and the flatworm, this gene is also expressed in the ciliary band of several other lophotrochozoan larvae (*13*). It is worth noting that ciliary cells are the only cell types in the two larvae that specifically share homologous TF expression.

Among neuronal clusters the strongest match is between oyster apical neurons and flatworm *MIP+* neurons, both of which are found in the apical organs of the larvae. A total of 23 orthologous genes are co-expressed in these clusters and among them the gene *Pde9* which is also found in the apical region of the larva of the annelid worm *P. dumerilii* (*48*) (see Fig. S10).

#### Comparison of lophotrochozoan larvae and echinoderm pluteus

We extended our comparisons to include data from the distantly related sea urchin pluteus larva (*8*). The only meaningful matches we recovered between oyster trochophore and pluteus larval cell clusters were ciliary cells and myocytes. Ciliary cells co-expressed a total of 116 oyster genes and 87 sea urchin genes. Among these we found many that overlapped with those shared between flatworm and oyster (*e*.*g*. such as: *annexins, calmodulins, calcineurins, centrins, cubulins, various dyneins, enkurin* and *tektins*). We found two TFs that were co-expressed (*Sp-FoxJ* & *Cg-FoxA/Cg-FoxL1* and *Sp-Pax9* & *Cg-Pax2/5*) however these appear to be paralogs and are not the same TFs as those shared between oyster and flatworm (for a full list see Fig. S11). SAMap analysis did not align oyster and sea urchin neurons nor skeletal cells to shell gland cells.

The comparison between the Müller’s larva and the echinoderm pluteus showed a good match between myocytes, neurons and gut cells. For pluteus neurons, the best match is with ciliary neurons of the flatworm larva; these share 20 flatworm and 16 sea urchin genes including several *neuronal acetylcholine receptors, calmodulins*, the gene *synaptogyrin* and *delta*. Delta expression is found in the brains of several other animals and in the apical neurons of the oyster larva (*16*).

We also find a match between the larval guts of Müller’s and pluteus larvae with cells co-expressing 116 flatworm and 61 urchin genes including several *dehydrogenases, glutathione S-transferase, ATP synthetases, glutamyl aminopeptidases, peroxiredoxin-6, retinol dehydrogenases, sterol carriers, apolipoproteins* and *heme binding proteins*. Pluteus and Müller’s larval guts also co-express the *macrophage mannose recepto*r gene (ManrC1A) which is a stomach specific marker of the sea urchin larval gut but do not co-express any TFs (*49*).

Finally, we observed a specific match between the exocrine pancreatic-like cell of the sea urchin and the gut-related cluster of the Müller’s larva. These clusters only co-expressed 19 flatworm genes and 17 urchin genes, however, many of these (*e*.*g. cpa2L, carboxypeptidase B, trypsin* and *pancreatic lipase-related* and *ptf1a* are very specific markers for the exocrine pancreatic-like cells of the sea urchin (*49*). Cells from these clusters are located in the upper gut in both larvae (although in the flatworm they are also found elsewhere around the gut) (Fig. 4). Exocrine pancreatic-like cells have been described in adults of cephalochordates, tunicates, cnidarians and in the pacific oyster (*50, 51*). Previously they have only been described in larval stages of the sea urchin *S. purpuratus* (*51*).

## General discussion

Ciliated larval stages are widespread across animals and their origins and evolution are longstanding problems in the field of evo-devo. Here, we present high-quality single cell atlases for larvae of two phyla within the super clade Lophotrochozoa: the trochophore larva of the oyster and the Müller’s larva of the polyclad flatworm. We used these datasets to explore the cellular composition of the two larvae in the contexts of their unique biologies and evolutionary histories and of their possible homology. We find that while the two larvae possess homologous cell types such as ciliary cells, neurons, myocytes and proliferative cells each larva presents its own idiosyncratic set of cells fitting it to its own biological niche.

The trochophore larva of the pacific oyster lacks several features that are commonly shared across lophotrochozoan larvae such as protonephridia, a fully differentiated gut and an apical tuft. Despite its fairly canonical trochophore-like appearance, the oyster larva also features several mollusc-specific innovations including a surprisingly large set of transcriptionally distinct muscle types and multiple cells involved in making the shell. The young transcriptomic age indices (TAIs) previously found for larval stages of bivalves using bulk transcriptomics are strongly affected by these clade-specific characters, most obviously by the derived molluscan shell transcriptome, and we suggest that measures based on bulk transcriptomics are not a reliable indicator of the recent evolution of trochophores (*31*).

The Müller’s larva is considered by some to be a highly derived trochophore and by others to be a convergently evolved novel larval type (*33, 52*). The Müller’s larva displays multiple classical larval characters missing in the mollusc trochophore, such as a large apical organ, paired protonephridia and a complex gut. These differences in larval complexity can be partially explained by alternative larval lifestyle: after the trochophore stage the oyster grows paired shells and quickly settles while the flatworm larva continues to swim in the plankton for over three weeks (*53*).

To test the homology of the similar cell types, we used SAMap to align our two datasets to each other and to the pluteus larva of an echinoderm. We found clear overlaps in orthologous gene expression between myocytes, proliferative cells, neurons, and gut cells. These are likely to be homologous at the level of adult Metazoa rather than necessarily demonstrating larval homology (*54*). We have, however, identified potentially homologous larval cell types including ciliary cells, a subset of neurons of the apical organ, and the larval gut.

The ciliary cells between oyster and flatworm larvae share over a hundred homologous genes, among these several TFs (namely *Dbx, Gsc, Otx, Pbx, Zfhx* and *Pax6*) and the spiralian specific gene *lophotrochin. Lophotrochin* and *otx* are also found in ciliary bands of several other lophotrochozoan and bilaterian larvae (*4, 13, 44*–*47*). When broadening the comparison to the sea urchin larval ciliary bands we still recover around 100 co-expressed genes between oyster and sea urchin (although no TFs) but not with the flatworm larva. Perhaps more significant is the identification of ciliary neurons located in the flatworm ciliary band. While equivalent cells are not present in the mollusc trochophore, these ciliary band specific neurons have been described in other canonical trochophores such as that of the flatworm *Platynereis* (*40*).

Our data also support a shared origin for apical neuronal cells - SAMap aligns the apical cells of the mollusc trochophore larva with a subset of neurons in the apical organ of the flatworm. These apical neurons co-express several orthologous genes, including the gene *pde9* which is also found in the apical organ of the annelid larva of *P. dumerilii* (*48*). In the flatworm (but not in the oyster) these neurons also express the protostome specific proneuropeptide *MIP* whose expression is reminiscent of what is observed in the larvae of the two annelids *Platynereis dumerilii* and *Capitella teleta* (*37*). We also find several TFs (namely *awh, hbn, delta* and *zic*) expressed in both sea urchin and oyster apical neurons (*8, 16*). Sea urchin neurons match better to the flatworm non apical ciliary-motor neurons, with which they share 20 genes including *delta*. We think this match is more likely to be due to a broad signature of neurons – indicating the likely homology of neurons – rather than a sign of apical organ homology. In fact, these shared TFs are generally found in the anterior nervous system across bilaterian adults.

The larval guts of flatworm and sea urchin share over 50 genes including some specific markers such as *ManrC1A* (*49*). We find a particularly strong match between the exocrine pancreatic-like cells of the pluteus larvae and a cluster of gut related cells in the flatworm that, similar to the pluteus, are located at the top of the gut (*8*). This finding further confirms the presence of pancreatic-like cells in protostomes and is only the second occurrence in ciliated larvae (*50*).

The co-expression of conserved TFs has been proposed as a key determinant of cell types across species (*55*) however, most of the co-expressed genes that we recover in cell types across larvae are effector genes; we find few shared TFs aside from those in ciliary band cells. We do recover several cell-type-specific TFs, ruling out technical limitations. This poses the question of whether cell type classification and homology can be based on TFs only or whether we should start considering the complete set of expressed genes. For instance, it would be interesting to test whether clearly convergent cell types share many homologous effector genes and/or any TFs at all.

Our results show that these two larvae are combinations of ancient cell types (such as myocytes and neuronal cells) that may be present in both adults and larvae and hence be poor clues of larval homology and clade-specific innovations (such as the shell gland in the oyster). Morphological structures can also be lost, as for instance the complex apical organ and protonephridia absent from the oyster larva. Even so we find that ciliary band cells and apical neurons share several orthologous genes and we consider that these are likely to be homologous to each other and to those of annelid larvae. This is the first cell type specific molecular evidence of the possible homology of polyclad flatworm larvae to other lophotrochozoan larvae. Our demonstration that individual larval types have gained and lost cell types shows that we will need to sample considerably more widely to answer the question of larval homology.

## Materials and Methods

### Animal husbandry

*Crassostrea gigas* individuals, raised in farms in Salcott Creek Essex, UK, were bought during the spawning season (May to August 2018 and 2019) from Richard Haward’s Oysters in Borough Market, London, UK. In the laboratory, oysters were kept at 16 ℃ in running artificial sea water and fed three times a week with Spirulina powder and invertebrate food supplement. For spawning, male and female *Crassostrea gigas* were shucked, gametes were stripped and put in glass beakers containing artificial filtered sea water (AFSW). Eggs were left in AFSW for about 1 hour to improve synchronicity then a dilution of sperm was added. After 5 minutes the water was tipped onto a 20 μm filter mesh and fertilised eggs were washed several times to avoid polyspermy. Fertilised eggs were then collected from the mesh and placed in beakers of AFSW at either 20 °C or 25 °C in an incubator. Trochophore larvae were collected on a 20 μm mesh after 16 h (25 °C) or 20 h (20 °C).

*Prosthecereaus crozieri* adult specimens were collected in coastal mangrove areas in the Lower Florida Keys, USA in October 2019. In the laboratory, animals were kept at room temperature (∼21 °C) in plastic boxes filled with artificial seawater. The water was changed daily for the first two weeks and then once every 2-3 days. The animals cannot be fed in the laboratory and so they were kept starved. Whenever eggs were found they were placed in separate containers and daily checked for hatchling. Hatched larvae were transferred into a filter and washed several times in filtered sea water then relaxed in 7.14% MgCl2 * 6H2O.

### Cell dissociation

*Crassostrea gigas* trochophore larvae were collected on a 20 μm filter mesh and transferred to low binding tubes. Larvae were washed several times in Ca2+ Mg2+-free artificial sea water with the aid of a centrifuge. Then animals were moved in a 4×4 well and were left in the solution for 3-5 minutes. After this time 300 μL of 0.5% Pronase (Roche cat # 10165921001) and 1% sodium thioglycolate (Sigma T0632) in low Ca2+ Mg2+-free ASW seawater were added and the solution was gently pipetted up and down to mix. After 3 minutes 10 μL of 5 mg/mL Liberase (Roche, cat # 05401119001) were added. The solution was mixed by gently pipetting up and down then larvae were very gently manually triturated. The whole procedure never lasted more than 20’ to reduce cell mortality. 1-day old larvae of *P. crozieri* were collected in a 40 μm filter and washed several times with filtered ASW. After cleaning the larvae were washed several times with Ca2+ Mg2+-free artificial sea water to prepare for dissociation. Larvae were collected in the centre of the mesh and transferred to a plastic cell culture petri dish and most of the NoCaNoMg-ASW was removed by pipetting. 300 μL of 1:100 solution of Prot14 (3.5u/mg; Sigma P5147) in low Ca2+ Mg2+-free ASW previously activated at 37 ℃ for 1 h were added. The solution was pipetted gently for 5-10’ until most of the larvae were dissociated. After this time the larval gut (which remained undissociated) was collected with the pipette and gently triturated until most dissolved and then added to the remaining cells. The dissociation process usually lasted around 15 minutes. For both animals, after dissociation cells were washed several times in Ca2+ Mg2+-free artificial sea water, checked for viability using Fluorescein Diacetate (*e*.*g*., Sigma F7378) and Propidium Iodide (*e*.*g*. Sigma P4170) and only samples with over 80% of viable cells were used for downstream capture.

### ScRNA seq data analysis

About 30,000 cells per sample were loaded into 10x chips following manual instructions using the 10x Chromium controller and Chromium single cell 3’ Kit v2, v3 or v3.1 (Cat #120237, 10x Genomics, USA). Four single cell libraries per animal were sequenced on Illumina NextSeq500 using a 2×75 paired-end kit to assess quality with shallow sequencing. Oyster reads were mapped against the chromosome scale genome of *C. gigas* generated by Peñaloza and colleagues (9). Flatworm reads were mapped onto the *P. crozieri* genome generated by our group (56). After shallow sequencing output matrices were analysed in Seurat v4.0.1 R according to the Seurat scRNA-seq R package documentation (57) (details of the following analysis can be found in the zenodo folder). Cells with less than 200 genes, more than 20% of mitochondrial genes and more than 5000 UMI were discarded. For the flatworm the mitochondrial cutoff was set to 30%. At this point we selected the best quality libraries from each animal for downstream analysis: library Cg1 for the oyster and Pc3 and Pc4 for the flatworm (Fig. S1 for the oyster and S5 for flatworm).

From these libraries we recovered 9,314 cells for the oyster and 17,605 for the flatworm. We then normalised the datasets by dividing gene counts for each cell by the total counts for that cell and multiplied by 10,000, then natural log transformed using log1p. Subsequently the top 2,000 variable genes were found using the vst method, datasets were scaled and principal component (PCA) analysis was performed. Nearest Neighbor (SNN) graph was computed with 70 dimensions and initial resolution of 10 then clusters were combined with a script for merging similar clusters from (58). Briefly, we calculated the average expression profiles for each cluster, then normalised expression vectors were used to calculate Pearson correlations between each pairwise cluster combination. Their correlation was ranked from highest to lowest and then we used a Wilcoxon rank-sum test to calculate the number of differentially expressed genes between each pair. We merged all cluster pairs that differed by less than 20 differentially expressed genes with 2-fold. This process was performed iteratively on all cluster pairs and gave us 39 genetically distinct clusters in the oyster larva and 51 in the flatworm larva. To confirm that by only selecting the highest quality libraries we had not missed any cluster we also similarly analysed all libraries per species together. Oyster libraries had to be integrated as described in (59). In this case we used a final resolution of 1 and results from this analysis can be seen in Fig. S1 and S5.

### Whole mount in situ hybridization/HCR

Whole embryo chromogenic in situ hybridisations for *C. gigas* were carried out as previously described (Osborne et. al 2018) in 96-well “U” bottom plate. Images were taken using a Zeiss Axio Imager M1. Probes for HCR were designed using the probe generator devised by Ryan Null (https://github.com/rwnull/insitu_probe_generator) and then ordered from IDT (Integrated DNA technologies), amplifiers and solutions were bought from Molecular Instruments. Samples were re-hydrated into PTW-DEPC (0.1% Tween-20 in 1x PBS), pre-hybridized in 200μL of hybridization buffer for 30’ at 37 °C and then 50μL of hybridisation buffer with 8nM each of probe were added. Samples were incubated overnight at 37 °C in a thermoblock shaking at 750rpm. The following day samples were washed four times in 0.5 mL HCR probe wash solution for 10’ at 37 °C then three times in 1mL 5XSSCT(DEPC) for 5’ at room temperature (RT). Samples were pre-amplified in 100μL of amplification buffer for 30’ at RT. At this point 2μL of each hairpin (for three probes experiments B1-H1, B1-H2, B2-H1, B2-H2, B3-H1, B3-H2) per experiment were placed in different PCR tubes. Heated in a PCR Thermocycler for 1:30’ at 95 °C, quickly spun down, and let it cool at RT for 30’ in the dark. Then all hairpins were pooled in one tube with 50 μL per experiment of amplification buffer. Then 50μL of amplification buffer and hairpin mix was added to all tubes with the samples (final concentration of 40nM Hairpin). Samples were incubated overnight at 25 °C in a thermomixer shaking at 750 rpm. The following day samples were washed three times in 1 ml 5X SSCT for 10’ at RT, then stained with DAPI (final conc. = 5 μg/ml) in 500 μL PTW for 15’. They were washed again twice in 500μL PTW and then transferred to 2,2ʹ-Thiodiethanol for imaging. The amplifiers used for the HCR experiments were B1-647, B2-594 and B3-488 and they were imaged using a Zeiss LSM-800 confocal microscope.

### Phylostratigraphy analysis

Gene age was estimated for both species using the GenERA tool (60). Briefly the tool implements genomic phylostratigraphy (61) by searching for homologs throughout the entire NR database and combining these results with the NCBI Taxonomy to assign an origin to each gene and gene family in a query species. Once we assigned each gene to its phylostrata we used the package myTAI to calculate the transcriptomic age index of the log transformed gene average expression per cluster, calculated using the AverageExpression function of Seurat.

Phylostrata enrichment analyses were computed using a hypergeometric test applied to the number of marker genes in each cluster belonging to each phylostrata compared to the global set of expressed genes.

### Cross-species cell-type comparison

SAMap v1.0.2 was used to compare the *C. gigas* and *P. crozieri* scRNA-seq datasets to each other, to the larval scRNAseq dataset of *S. purpuratus* (8) and to the adult scRNAseq dataset of *S. mediterranea* (32).

### Neuropeptide precursor search

Neuropeptide precursors were identified by two different methodologies. First, BlastP analyses were performed using previously published neuropeptide datasets with an e-value of 1e-2 (37, 62–64). Then, predicted secretomes for *C. gigas* and *P. crozieri* were obtained using SignalP4.1 with the sensitive option (D-cutoff 0.34). This secretome was used to search for novel precursors by regular expressions based on dibasic and monobasic cleavage sites as described before (64). These two methodologies produced a large database that included hundreds of hits that were manually curated by comparing with the neuropeptide precursor complement from several lophotrochozoan species (37, 64) (see Fig. S11, S2 and S13).

## Acknowledgments

We would like to thank Johannes Girstmair and the Florida Keys Marine Lab for their help with animal collection, Alexander Tarashansky for his help with running SAMap and Kate Rawlinson for her advice on ISH in the polyclad flatworm and for providing us the sequence for *r-opsin*.

