## Supplementary Materials for "Single-cell atlases of two lophotrochozoan larvae highlight their complex evolutionary histories"

### Supplementary material

**Fig. S1 Quality assessment of initial shallow sequencing of oyster trochophore scRNA libraries.** A) Violin Plots showing gene number per cell (nFeature_RNA), UMI per cell (nCount_RNA) and percentage of mitochondrial genes (percent_mito) per cell in each sample (Cg1, Cg2, Cg3 and Cg4). Cg2 and Cg3 are technical replicates from the same dissociation. Sample Cg1 (used for downstream analysis) presents more overall cells, higher genes and UMIs and lower mitochondrial gene content. B) UMAP of integrated samples Cg1, Cg2, Cg3 and Cg4 coloured by cell clusters C) UMAP of integrated samples Cg1, Cg2, Cg3 and Cg4 coloured by sample of origin shows Cg1 cells are present in all clusters.

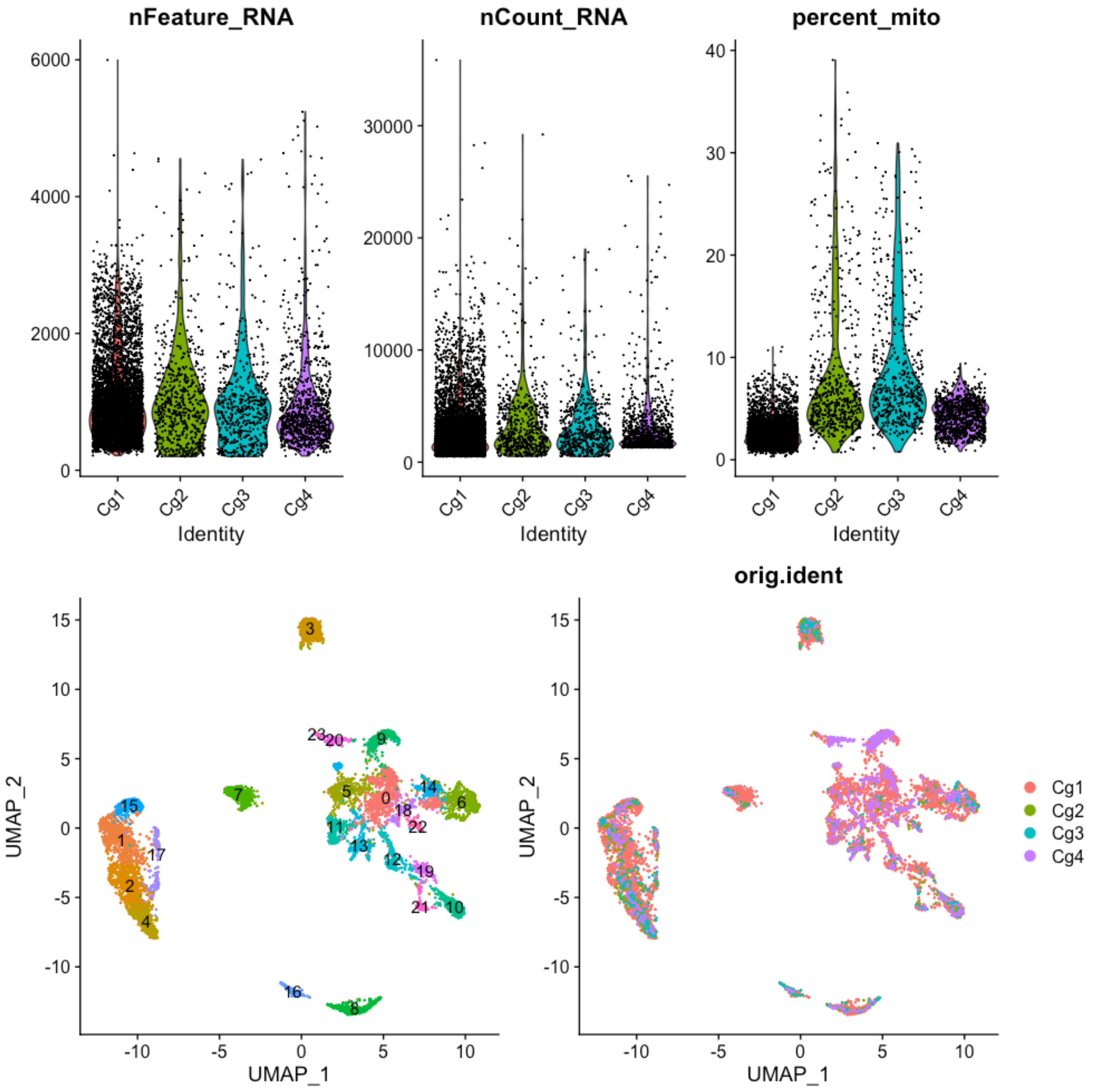

**Fig. S2 Several lophotrochozoan specific genes show expression in the oyster larva ciliary band clusters.** Dotplots show expression of genes (x axis) in each cell cluster (y axis) of the oyster scRNAseq, blue dots indicate average expression, size of dots indicate percentage of cells expressing the gene. Genes shown here are Lophotrochozoan specific genes from a study by Wu and colleagues (*13*).

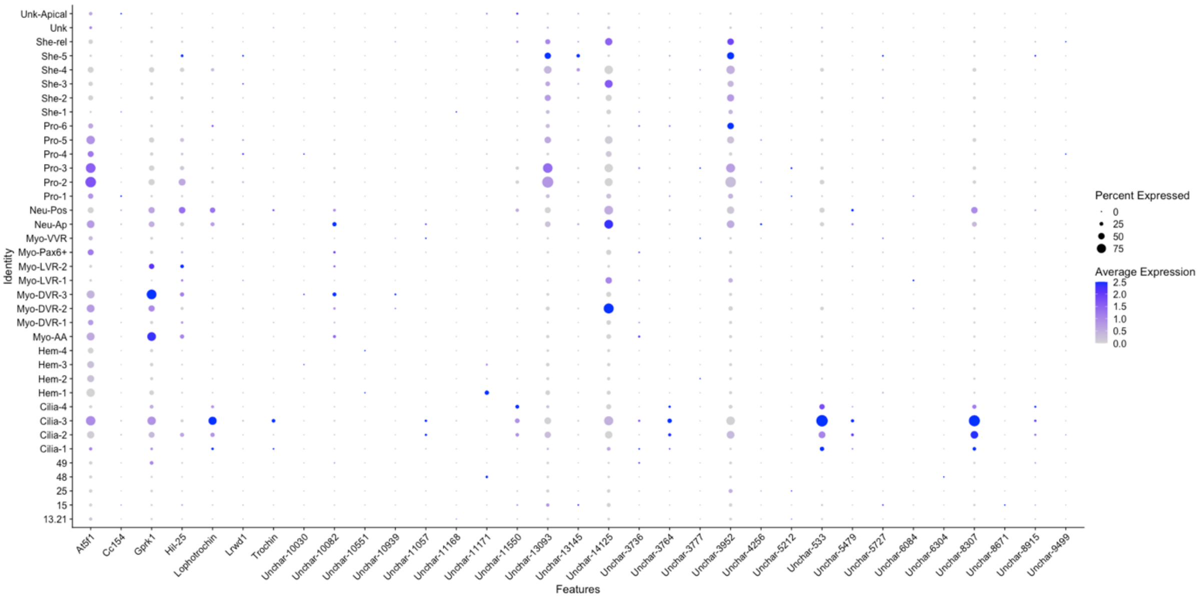


**Fig. S3 Neuropeptides and neuronal genes expression in the trochophore (left) and Müller's larva (right) neuronal clusters.** Dotplots show expression of genes (x axis) in each cell cluster (y axis) of the *C. gigas* scRNAseq (left) and *P. crozieri* (right), blue dots indicate average expression, size of dots indicate percentage of cells expressing the gene. Genes shown here are a selection of neuronal markers and neuropeptides present in both animals.

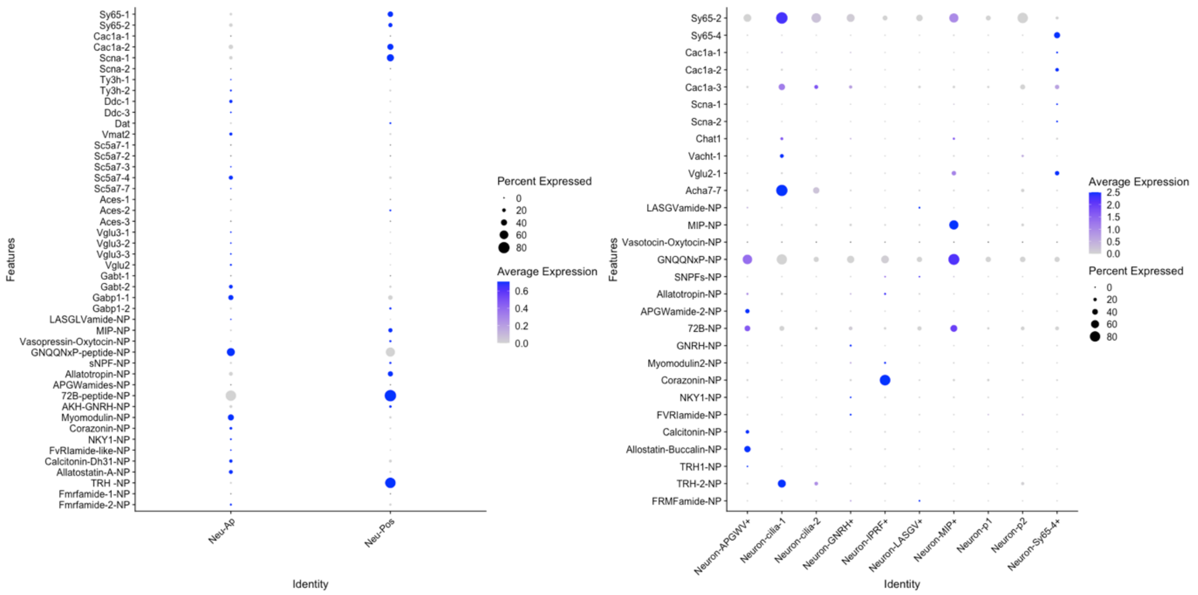

**Fig. S4 Expression of TFs in the Oyster larva scRNA. Different myocytes clusters express different subsets of TFs.** Dotplots show expression of genes (x axis) in each cell cluster (y axis) of the oyster scRNAseq, blue dots indicate average expression, size of dots indicate percentage of cells expressing the gene. Genes on the x axis are all TFs found in the oyster larva that are markers for a cluster.

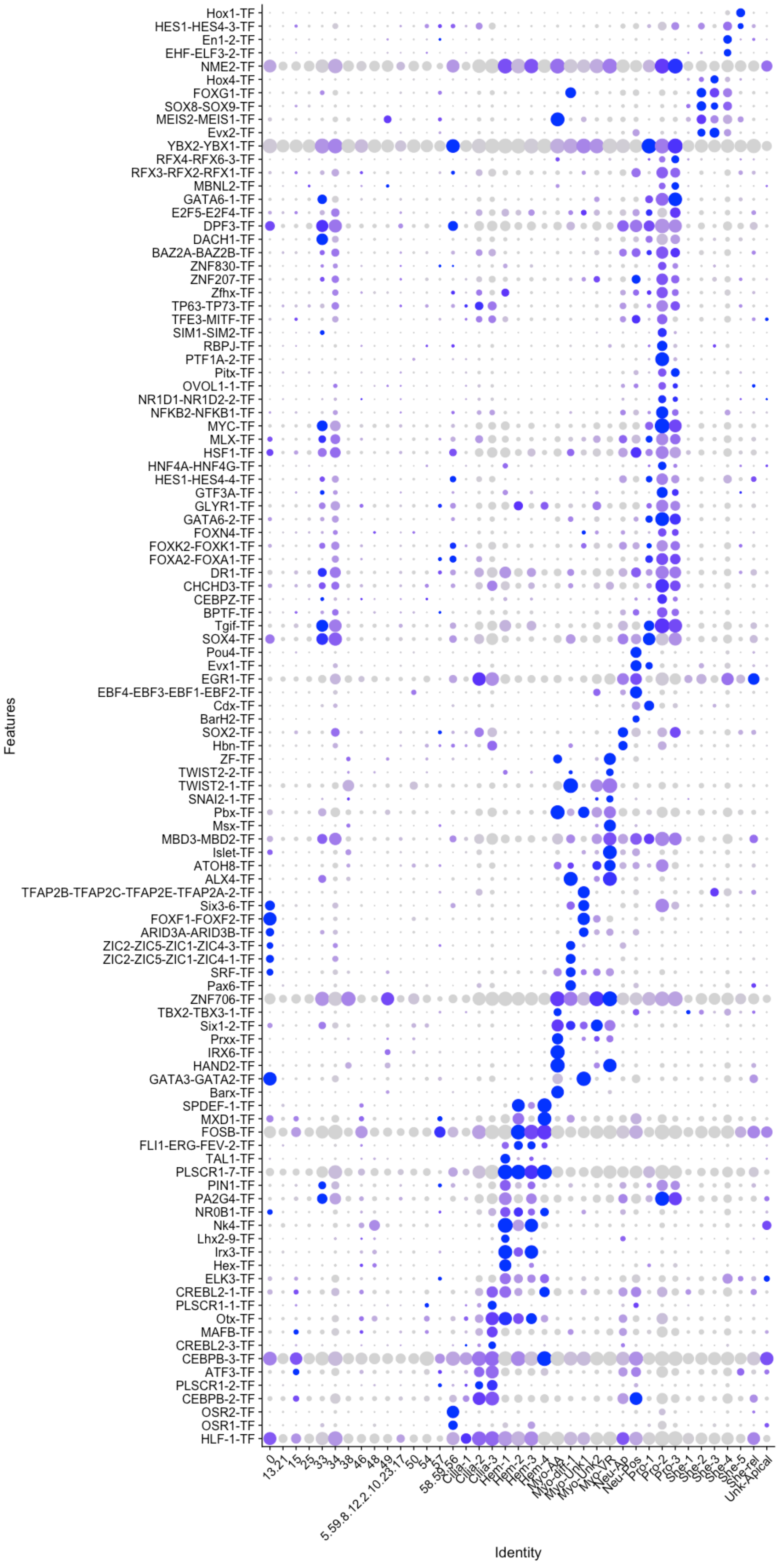
Hox1-TF
HES1-HES4-3-TF
En1-2-TF
EHF-ELF3-2-TF
NME2-TF
Hox4-TF
FOXG1-TF
SOX8-SOX9-TF
MEIS2-MEIS1-TF
Evx2-TF
YBX2-YBX1-TF
RFX4-RFX6-3-TF
RFX3-RFX2-RFX1-TF
MBNL2-TF
GATA6-1-TF
E2F5-E2F4-TF
DPF3-TF
DACH1-TF
BAZ2A-BAZ2B-TF
ZNF830-TF
ZNF207-TF
Zfhx-TF
TP63-TP73-TF
TFE3-MITF-TF
SIM1-SIM2-TF
RBPJ-TF
PTF1A-2-TF
Pitx-TF
OVOL1-1-TF
NR1D1-NR1D2-2-TF
NFKB2-NFKB1-TF
MYC-TF
MLX-TF
HSF1-TF
HNF4A-HNF4G-TF
HES1-HES4-4-TF
GTF3A-TF
GLYR1-TF
GATA6-2-TF
FOXN4-TF
FOXK2-FOXK1-TF
FOXA2-FOXA1-TF
DR1-TF
CHCHD3-TF
CEBPZ-TF
BPTF-TF
Tgif-TF
SOX4-TF
Pou4-TF
Evx1-TF
EGR1-TF
EBF4-EBF3-EBF1-EBF2-TF
Cdx-TF
BarH2-TF
SOX2-TF
Hbn-TF
ZF-TF
TWIST2-2-TF
TWIST2-1-TF
SNAI2-1-TF
Pbx-TF
Msx-TF
MBD3-MBD2-TF
Islet-TF
ATOH8-TF
ALX4-TF
TFAP2B-TFAP2C-TFAP2E-TFAP2A-2-TF
Six3-6-TF
FOXF1-FOXF2-TF
ARID3A-ARID3B-TF
ZIC2-ZIC5-ZIC1-ZIC4-3-TF
ZIC2-ZIC5-ZIC1-ZIC4-1-TF
SRF-TF
Pax6-TF
ZNF706-TF
TBX2-TBX3-1-TF
Six1-2-TF
Prxx-TF
IRX6-TF
HAND2-TF
GATA3-GATA2-TF
Barx-TF
SPDEF-1-TF
MXD1-TF
FOSB-TF
FLI1-ERG-FEV-2-TF
TAL1-TF
PLSCR1-7-TF
PIN1-TF
PA2G4-TF
NR0B1-TF
Nk4-TF
Lhx2-9-TF
Irx3-TF
Hex-TF
ELK3-TF
CREBL2-1-TF
PLSCR1-1-TF
Otx-TF
MAFB-TF
CREBL2-3-TF
CEBPB-3-TF
ATF3-TF
PLSCR1-2-TF
CEBPB-2-TF
OSR2-TF
OSR1-TF
HLF-1-TF
Features
0
13.21
15
25
33
34
38
46
48
49
5.59.8.12.2.10.23.17
50
54
57
58.53.56
Cilia-1
Cilia-2
Cilia-3
Hem-1
Hem-2
Hem-3
Hem-4
Myo-AA
Myo-diff-1
Myo-Unk1
Myo-Unk2
Myo-VR
Neu-Ap
Neu-Pos
Pro-1
Pro-2
Pro-3
She-1
She-2
She-3
She-4
She-5
She-rel
Unk-Apical
Identity

**Fig. S5 Quality assessment of initial shallow sequencing of the flatworm Müller's larva scRNA libraries.** A) Violin Plots showing gene number per cell (nFeature_RNA), UMI per cell (nCount_RNA) and percentage of mitochondrial genes (percent_mito) per cell in each sample (Pc1, Pc2, Pc3 and Pc4). Sample Pc3 and Pc4 (technical replicates used for downstream analysis) present more cells, higher genes and UMIs and lower mitochondrial gene content. Pc1 and Pc2 are technical replicates of each other from the same dissociation. B) UMAP of integrated samples Pc1, Pc2, Pc3 and Pc4 coloured by cell clusters C) UMAP of integrated samples Pc1, Pc2, Pc3 and Pc4 coloured by sample of origin shows that cells from Pc3 and Pc4 libraries are present in all clusters.

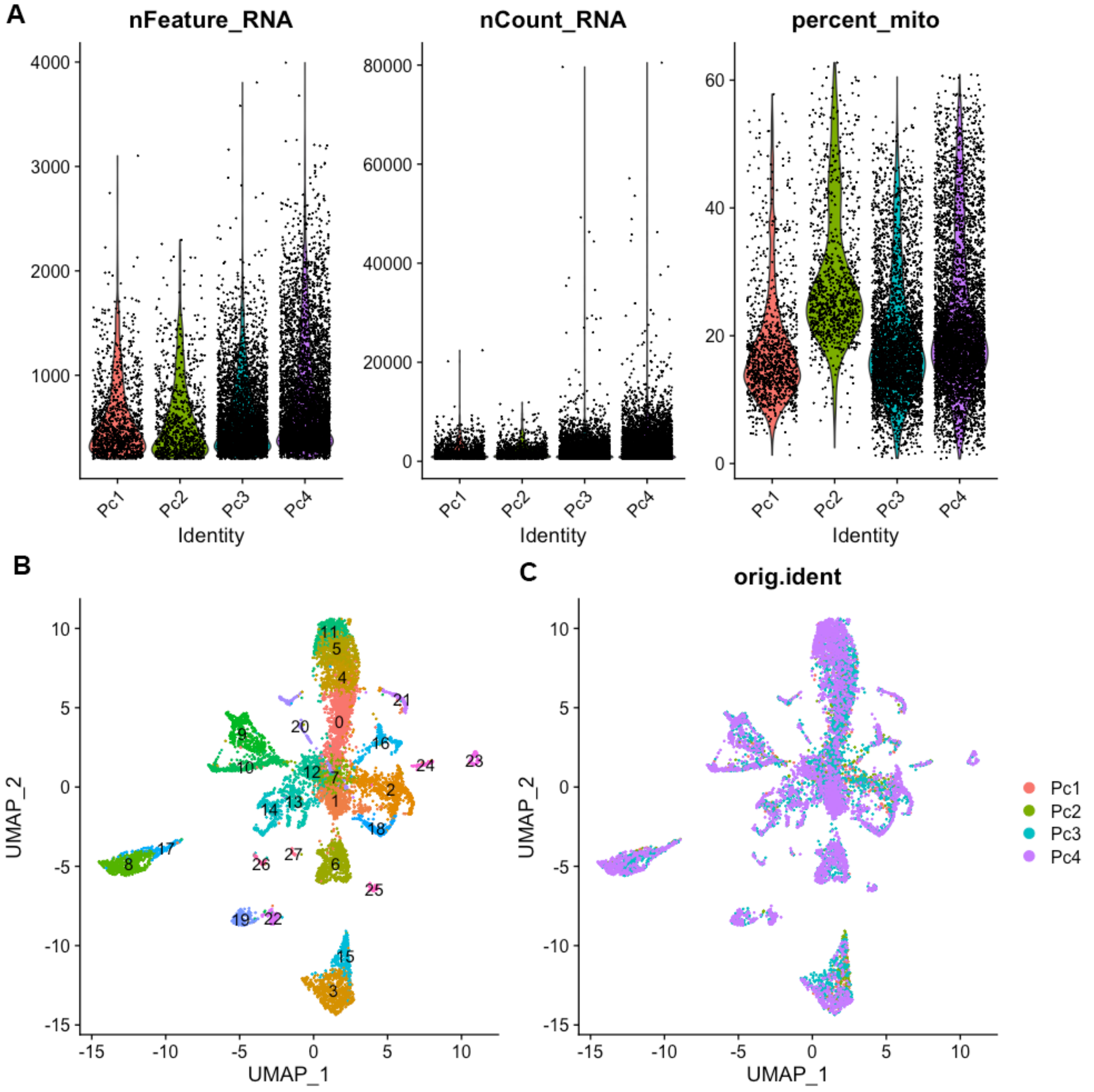

**Fig. S6. SAMap alignment between S. mediterranea adult and P.crozieri larvae show similarities between cathepsin cells, gut cells, myocytes, neurons and ciliated cells.** *S. mediterranea* scRNAseq used here is from (*32*). SAMap alignment scores are defined as the average number of mutual nearest cross-species neighbors of each cell relative to the maximum possible number of neighbors (*42*).

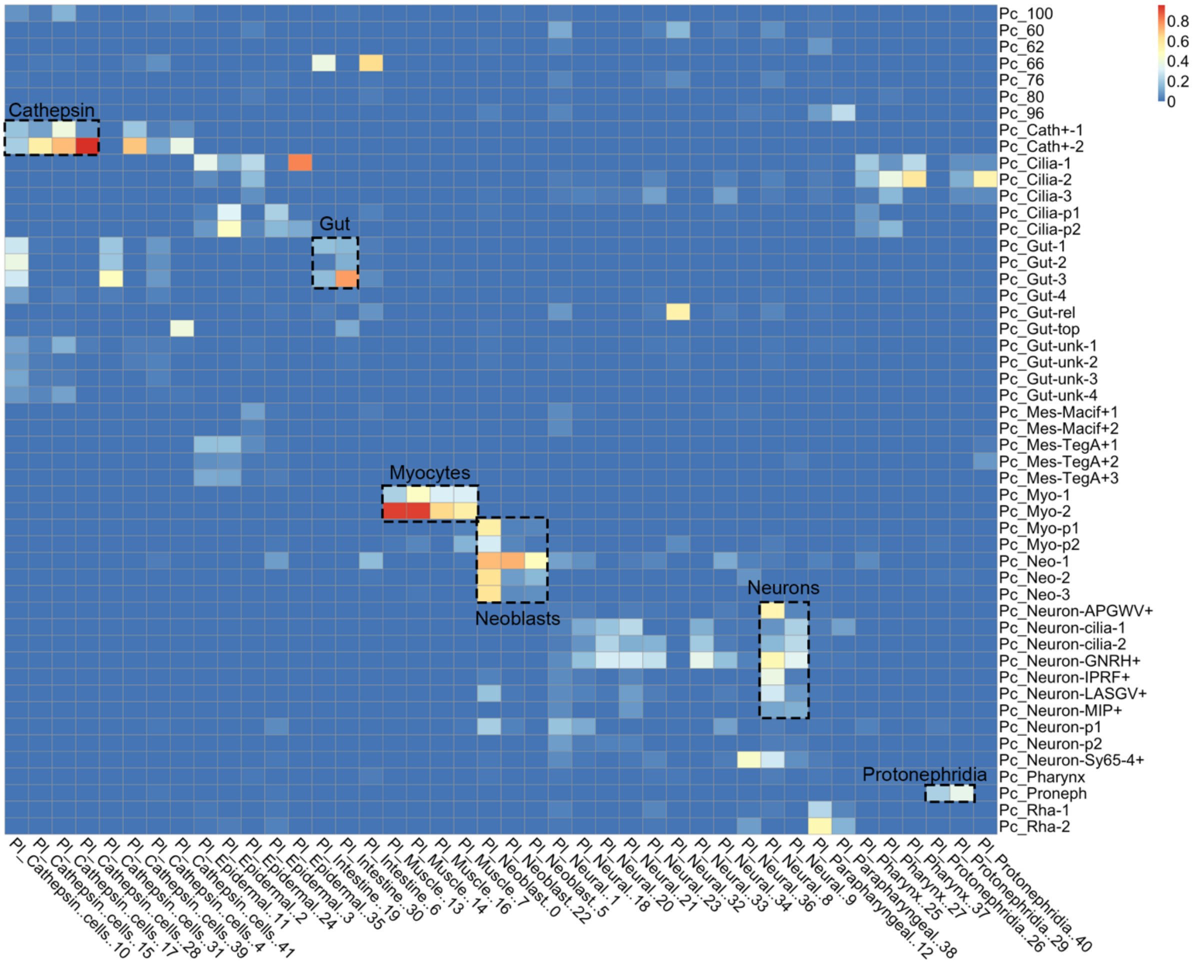

**Fig. S7. Gene age analyses in different cell types of the flatworm larva.** A) Transcription age index (TAI) for different cell types, smaller TAI values correspond to “older” gene age. Gene age is inferred using a phylostratigraphy approach, then transcriptomic age index is calculated on the log transformed gene average expression per cluster. B) Phylostrata enrichment analyses per cell type. Phylostrata enrichment was computed using an hypergeometric test applied to the number of marker genes in each cluster per phylostrata compared to the global set of expressed genes.

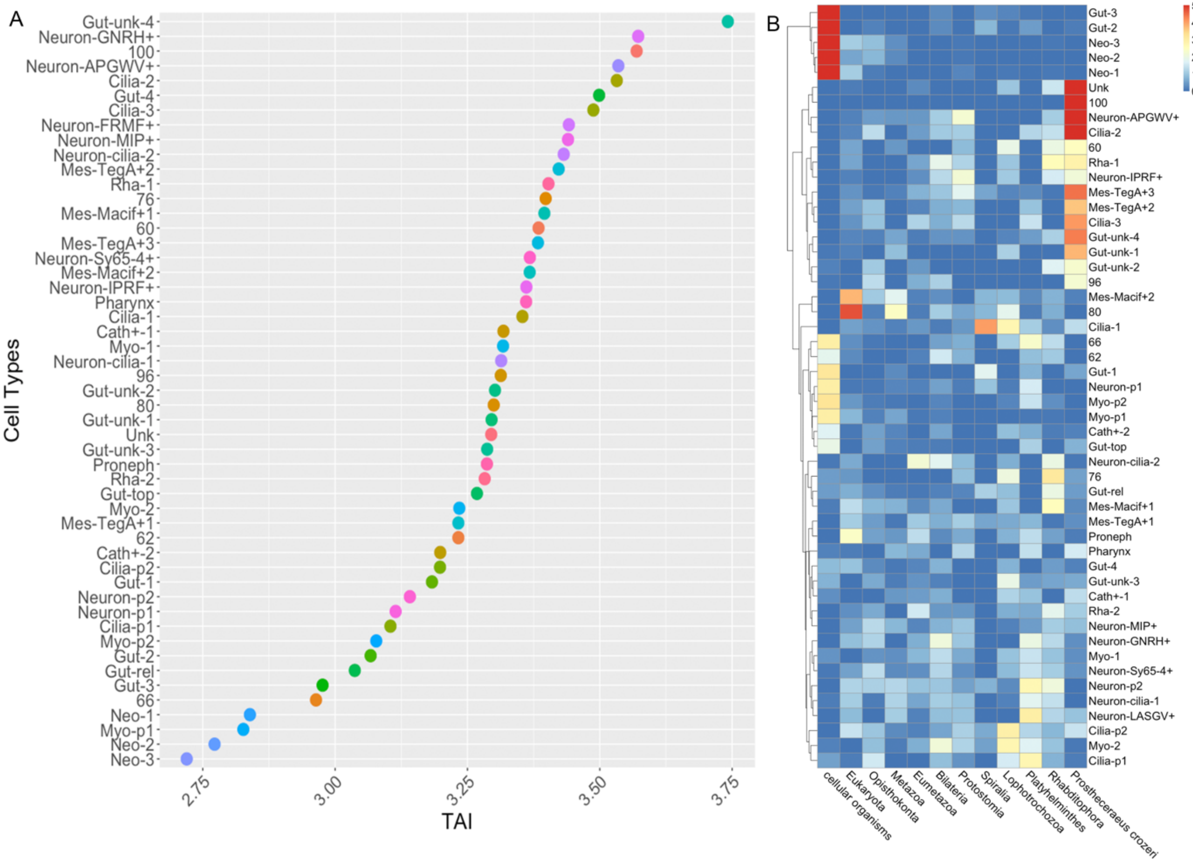

**Fig. S8. Ciliary genes co-expressed between Oyster and Flatworm larva generated with SAMap.** SAMap calculates genes that are co-expressed between each aligned pair of cell types. Here we show the expression of co-expressed genes between the oyster (left) and the flatworm (right) ciliary clusters. Dotplots show expression of genes (y axis) in each cell cluster (x axis), blue dots indicate average expression, size of dots indicate percentage of cells expressing the gene.

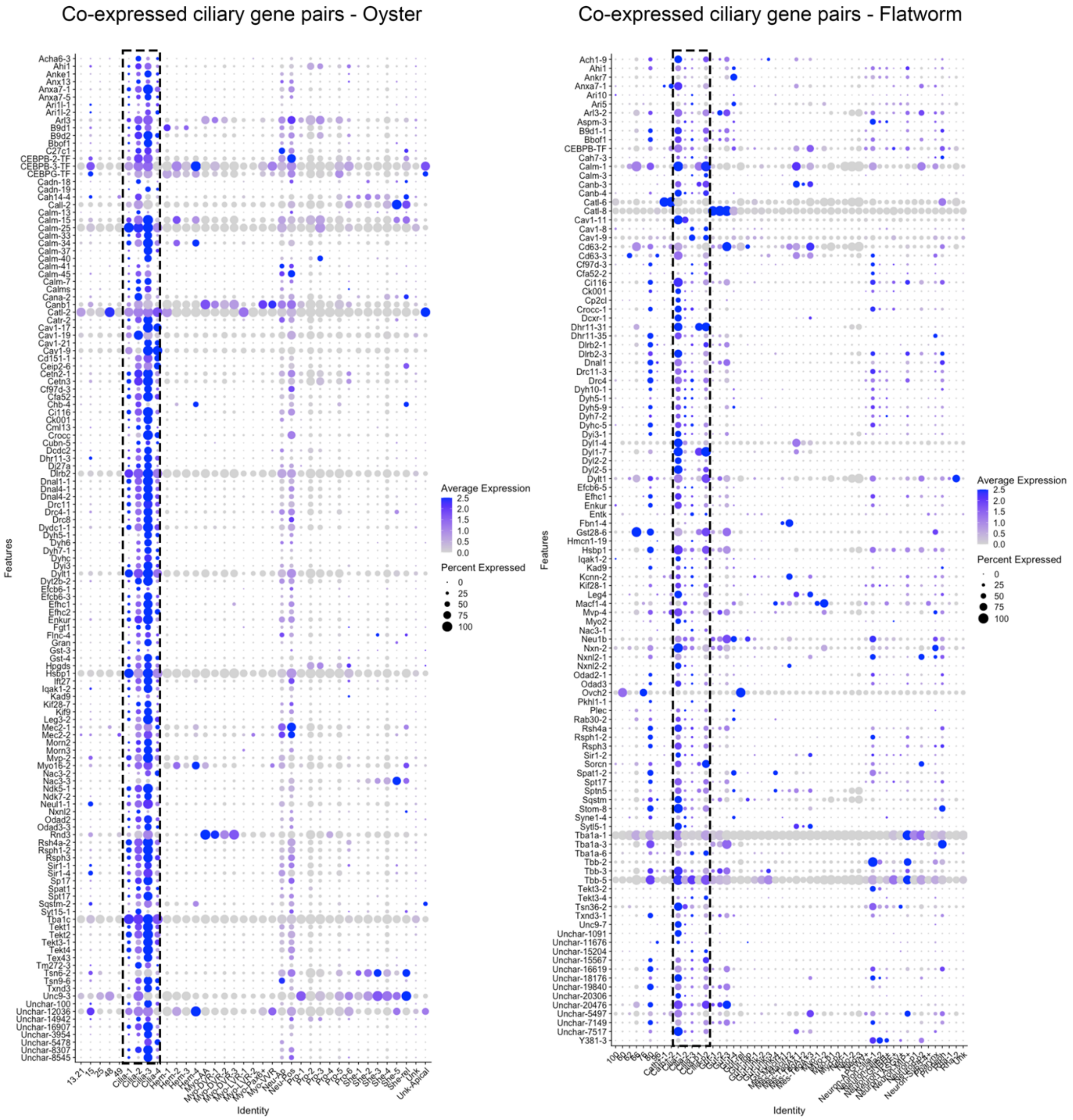

**Fig. S9. Expression of homologous TFs in ciliary bands of the oyster (left) and flatworm (right).** Dotplots show expression of genes (x axis) in each cell cluster (y axis) of the oyster scRNAseq (left) and flatworm (right), blue dots indicate average expression, size of dots indicate percentage of cells expressing the gene. Genes on the x axis are TFs expressed in the ciliary bands of both animals.

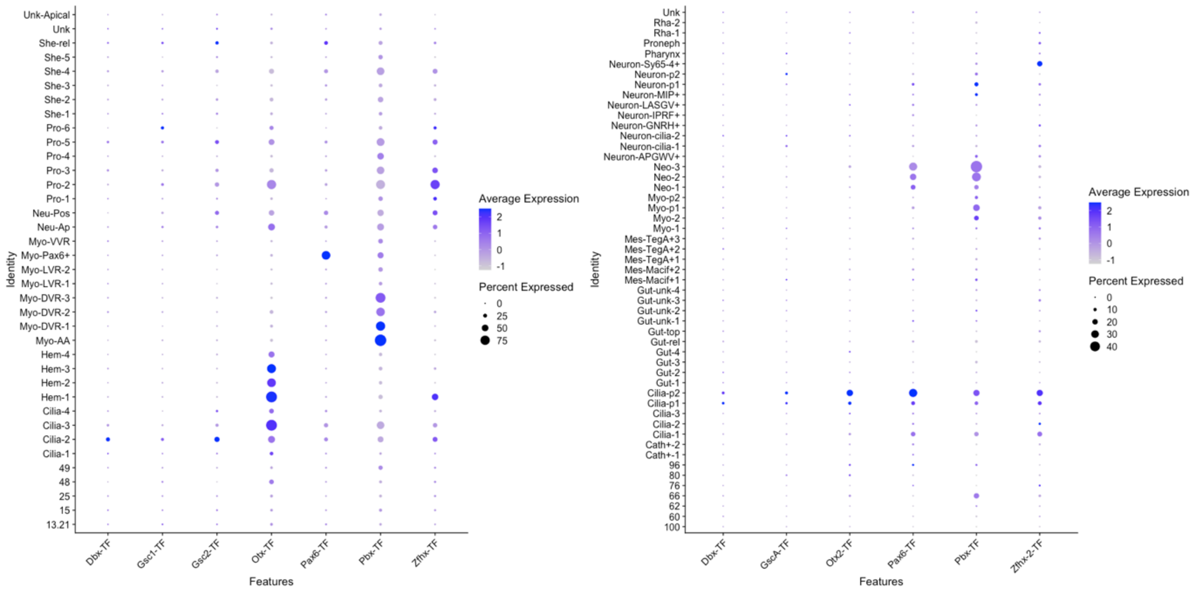


**Fig. S10. Co-expressed genes between the apical neurons of the oyster (left) and the MIP+ neurons of the flatworm (right).** SAMap calculates genes that are co-expressed between each aligned pair of cell types. Here we show the expression of genes co-expressed between the oyster apical neurons (left) and the flatworm MIP+ cells (right). Dotplots show expression of genes (x axis) in each cell cluster (y axis), blue dots indicate average expression, size of dots indicate percentage of cells expressing the gene.

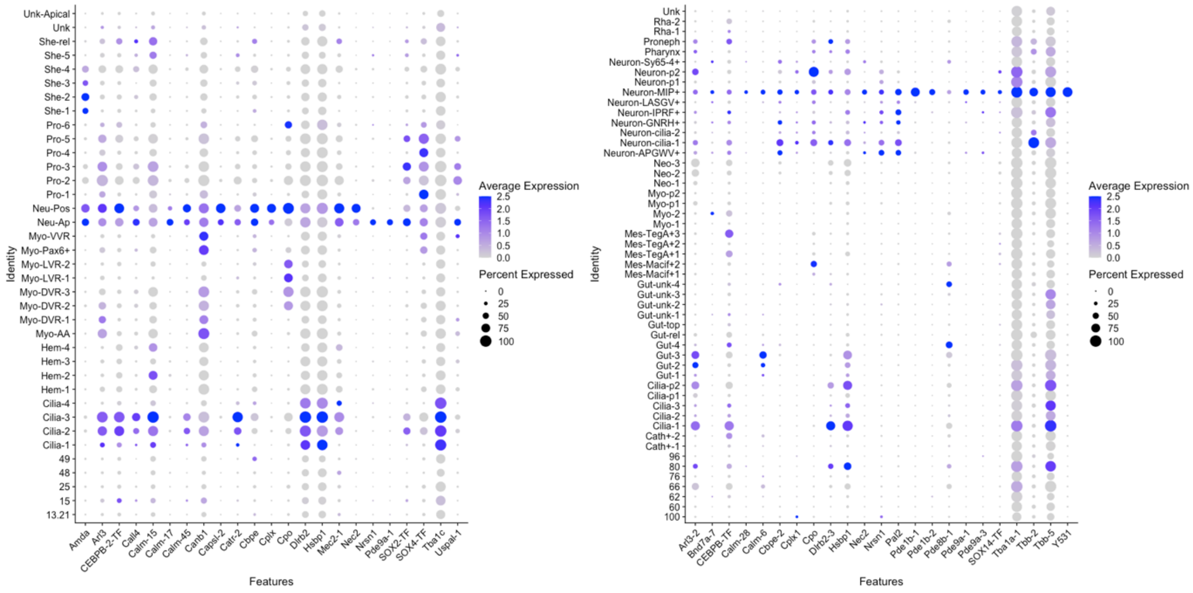

**Fig. S11 Ciliary genes co-expressed between Oyster and Sea Urchin larva generated with SAMap.** SAMap calculates genes that are co-expressed between each aligned pair of cell types. Here we show the expression of genes co-expressed between the oyster (left) and the flatworm (right) ciliary clusters. Dotplots show expression of genes (y axis) in each cell cluster (x axis), blue dots indicate average expression, size of dots indicate percentage of cells expressing the gene.

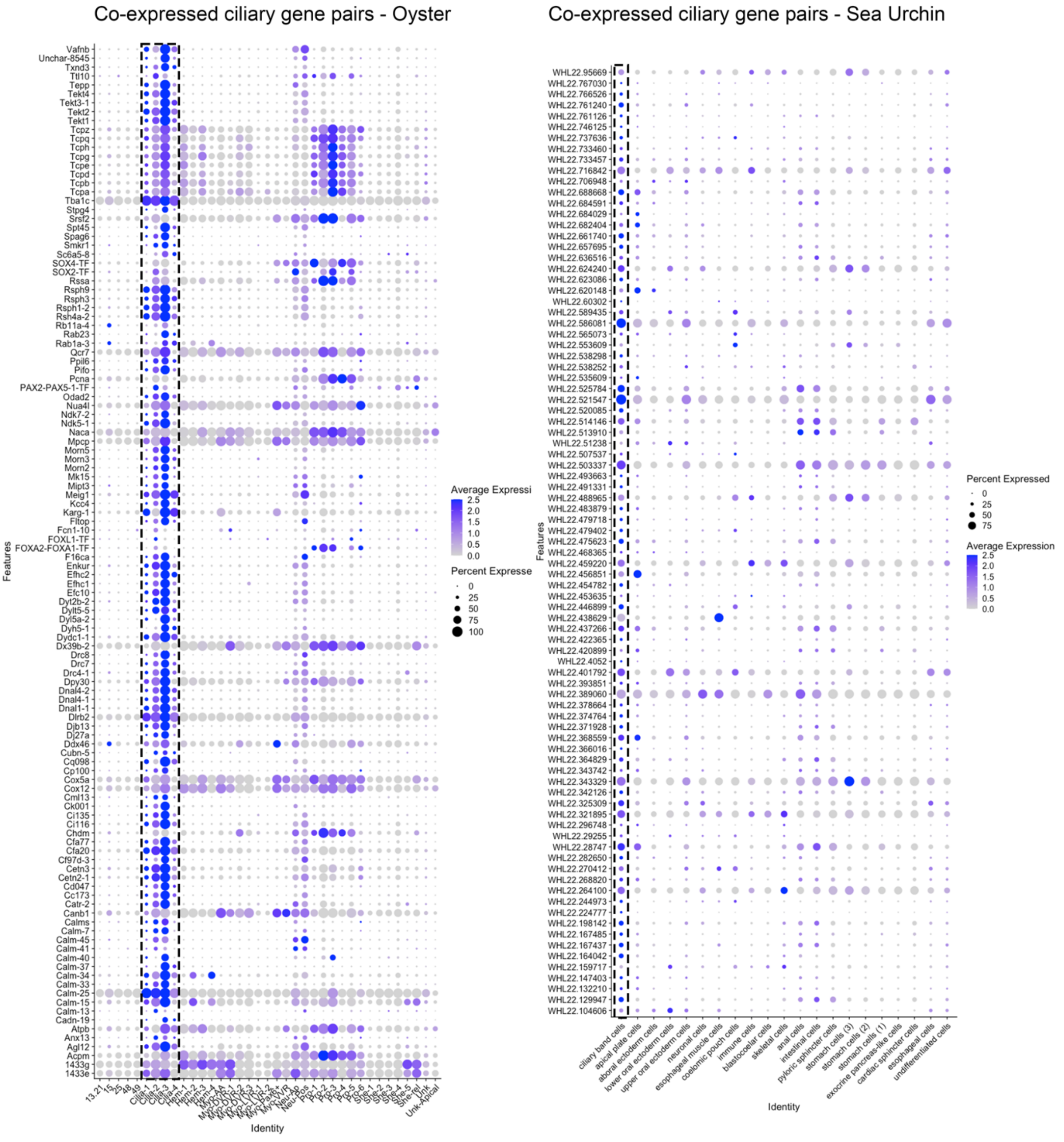

**Table S1. Presence (navy) and absence (white) of a set of Neuropeptides in Lophotrochozoa.** The original data for brachiopods, nemertans, *P. dumerilii* and phoronids is from (64).

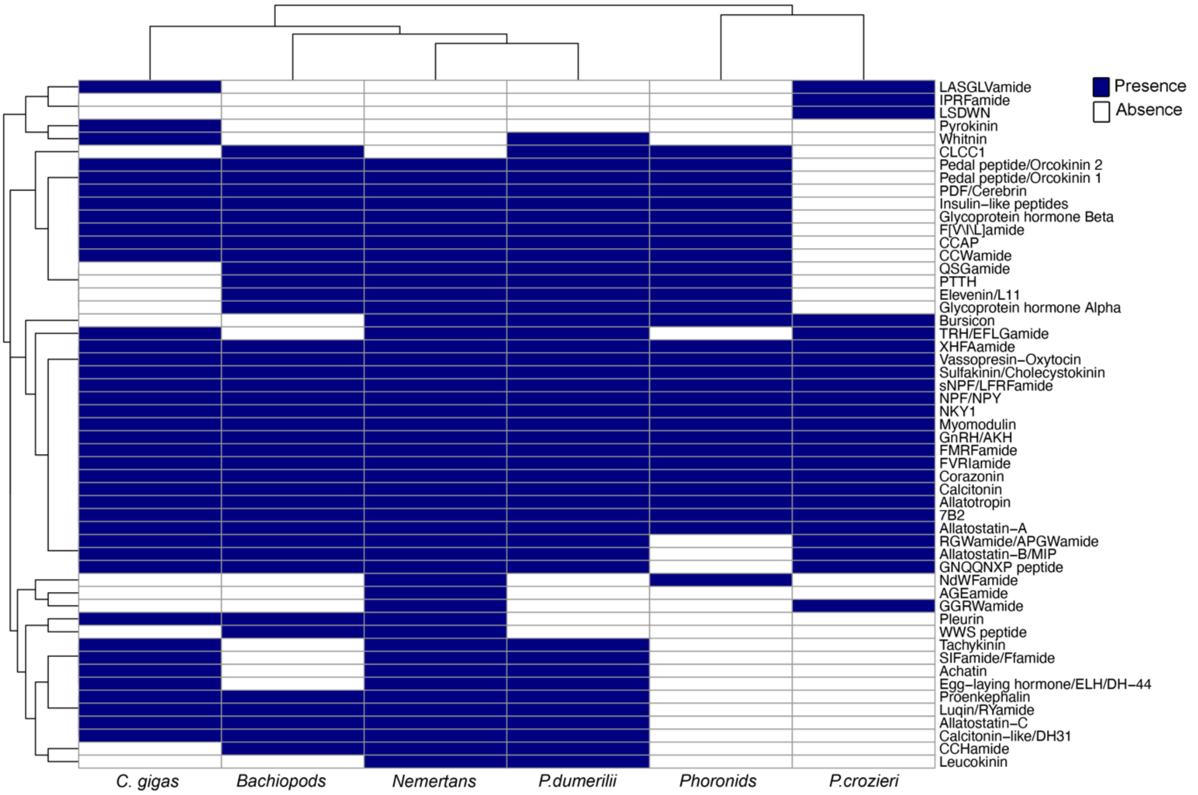


**Table S2. Sequences of *Crassostrea gigas* neuropeptide precursors.** A total of 47 neuropeptide precursors were identified in this species. The predicted signal peptide is shown in blue, predicted monobasic/dibasic cleavage sites are shown in green and predicted mature neuropeptides are shown in red.

>Cgig_7B2 peptide precursor
MAFTMILSVLAISFLMAEAKVSPDGSLVWPDLGQLYALDDLRNGLGNGYYDLESRS
SPNQDWSNAGYQEPTLNDDIFSEASIRDQEYQENSPLWGYQSVSGGTGDGKDNP
KQVKTDKVLPAY**CNPPNPCP**IGYTSDDNCVENFENSPDNNRKLMKQQECPCDKEH
MFSCPSDTSISTKSGPSLDSVIEQLGNMENNAYMSTNEKRKSLVAKKSPHLVKREA
EQARNPYFDGPHNKVAAKKGSYIPRHQ*
>Cgig_Achatin
MRTPLLILLISLAFTFGTAENHLLEQYENDEDIQHQLQNEAYYKRGFWDKRGFWDK
RMSHKDFSHGSDDSSWNNLILLLKDYIRYKHESNKKEQVFSNGLQNSH*
>Cgig_Allatostatin-A
MWSTNYATTVFGFFCFVQVFVLTVSQHISSHNDDYTKHLENIKFLNKEAEISPKQPA
DDDVDFGNSDTADDLSDLTEEEKRALDRYSFSGSLGKRGLDRYSFYGGLGKRALD
RYGFFGGLGKRALDQYGFAGSLGKRALDRYSFMGGLGKRKLDQYGFAGRLGKRA
LDRYGFVGTLGKRKLDQYSFMGNLGKRRLDSHRYFGSLGKRALDRYGFFGGLGK
RADTLGNSQENIQGADKDEKFEQKRYGFFGGLGKRALDQYGFAGSLGKRALDRYS
FMGGLGKRKLDQYGFAGRLGKRALDRYGFVGTLGKRKLDQYSFMGNLGKRRLDS
HRYFGSLGKRALDRYGFFGGLGKRADTLGNSQENIQGADKNEKFEQKRLYPYWY
YRQGGSPIYTQTRGIDRFSFAARLGRR*

>Cgig_MIP/Allatostatin-B Wamides
MLCHLEQLLLSCIVLCLVKVCASLKAQDEASVNDHDIVRRQAMGHSFGDELEGFDL
DPKRVNNWNQFPAWGKRLSKRRWSSLGAWGKRSWLDRLISANNNWGKRWKSM
SNSWGKRQAPSEFDGLSDDYINIKKRSVDSKFSHNSRNKRSIPTELSPEQNEEKRR
WSSLSAWGKRSDDDEKRRWSSLSAWGKRSNPEAIDDNDSDNISKRKWSSFSSW
GKRGDPVDLSKRLYSYWQNRLMTNNPWMERRGWNAFSSWGKRSMD
>Cgig_Somatostatin/AllatostatinC
MELTQSVFVLKLYAAVVAVLLVAEVHAQPQKFSTEIQQTGDESSTDLNLFKMALRE
AYNRELEFYEQQEAQIVKQLAALENDRNQIRERKRSHIRCLVNVIACYRKK*
>Cgig_Allatotropin
MQMVKVVLVTVLLTLCVIVDCLPHSSTKMRQKRGFRQSIVDRMGHGFGKRANTDL
YDPYLTKTPNLMTADELTTNILNSEDLAQAIVKKFIDLDGDGIISTAELLRTTS*
>Cgig_Calcitonin-like
MEGLKTSICSLLFFIVATTSLEIKSQHIREKHLVDQLDKVREQLKDLSHSVDTLRTFT
QREACALSLNVDICTEKYIEETADHQSKLQNLIEGNPGKRRPKRAADMSLDLFVEDL
LRKNAALQSIQQILNNMSKTVHQEKKRSCTLNLAYHCQTSEYAGLTDLYNYLNSNA
SPGKRR*
>Cgig_Calcitonin/DH31
MIAAFATDTRHIRSAISSAEEVKRETQEEQIKLCRGMGPNNHPCGLTSFDVRNVRR
GDDGDIEVETLNRGTRSLEGSRPDNPPTEKERMAYIAKLAKRRQDMDRILELLNEK
ENNIRQIRKRTCAVELGGACRTEWASAIADQYYYLMGPHGPGKRRRRSLINVLTGK
LSHISTSNKGH*
>Cgig_CCAP peptide partial
QNENSKLGVKKLAALLANELLSEDKDITQKDKRVFCNGFFGCSNGKKRSNSMNLN
PLDYPVDAPEPETKTDFRKRLFCNTGGCFGRRKRSTSQTLERRLNNRGSKDGNQ*
>Cgig_CCWamide
MTYFIAGIWILLISLNFGSCMYYADYDDDYYNQNKALDLFKSMDRRNYYLRQPICNS
WNRHCLPWSNEIQHSCCGGLACKCNLWGQNCRCTTQLWGR*
>Cgig_Corazonin
MKVSPCTQVIVMVLTLGLLCEVHAQNYHFSNGWQPGKRSYRGCTVRPEIRSILFKII
EDEVERIQKCSHSNIEDVFSLIQEKTGVDAREV*
>Cgig_Egg laying hormone_like
MKLNVMAVIVTALFVSVDAFILQDQDNQPDLSEADSIPVEKRGRLSLTADLRSLARM
LEAHRKRFIASRFPYDSIRKKLFRYGKRSPVPETLEYNEGEDRSNALDSEFPSLKVE
EESPIYNIKRQRLSVNGALSSLADMLAANGRQRMMSEMAMNRQRLFGLGK*
>Cgig_LRNFVamide-like_F[V/I/L]amide_like
MRVPVFLYSFVVFLGVCCVDEVISTDKDIESSSQEIDHQRDFATLNKRRYGDDDFQ
KRLRYFIGKRSDTGDKEKRMRYLLGKRFAFDGNVNKRLRYFLGKRDFSDTKKRTR
YFIEKRSDELGSHDLEQNMRHILEKQESDKVNPGLEKRMRYFLGKRMRYFLGKRA
NSATSNSDSQNESSKRTRYFLGKRDGTRYFLGKRFRYFLG*
>Cgig_FVRIamide-like
MLRPYHVIIVGLFYCYTTNAEINENKLLHPIKTEEGANEILGDKADDKRSRGFFRIGK
KSAVENEANDKKFDSKTVKEEDNYIPEKIQLIRVVNSESETPIYVPVEFDPESSDDTA
DEDEKRASGFFRIGKSAENVDKRKGFFRIGKSVDQNPMNKKASGFFRIGRTPIDKR
GKGFFRIGKSLNEMDEKRASGFFRIGKSALNDKRSRGFFRIGRSKGFFRIGKAFPLE
GEKRASGFFRIGRNSPEEMRKKASKFFRIGKSVNSKEENDKRASGFFRIGKKCSGD
SDKAGDNLTEDKSQSNPENEDTSESFNQNSDEPVRRASQFFRIGKSSSNKVTKRS
SGVNSSPEQNLNLNKRAFFRIGKVPTSAFMRIGRQHLLQSLVSDPLYRNGRIQQSS
FIRIGKRSMSDNHLIDDEQSDSSM*
>Cgig_FMRFamide

MGTWTYLCLLVAFLLNWFTIETSANDLIDDCYRNPELCQEVGILFGQQQPVDKRFL
RFGKRALSGDHYIRFGRNSDDKRFLRFGKRGEQGSVEDDLREALNKVIKFKQETGL
HLRKRRSADPPLVKDVPEDKDSNSTEKEDSASEKHKRETDEVSEENKRFMRFGRT
PAEDDPTYMKRFMRFGRNPDLEKKFMRFGKDGNEKRFMRFGKREDNDNMVTDD
KRFMRFGRDPKDDTLMERFVRNGRSGDDKRFMRFGKRFMRFGKRFDNDGEYDD
EDEMGAEKRFMRFGKSGDEEKRFMRFGKSGDEEKRFMRFGKSGDEEKRFMRFG
KSGDDEKRFMRFGKSGDEEKRFMRFGKSGDEEKRFMRFGKSGDEEKRFMRFGK
SGDEEKRFMRFGKSGDEEKRFMRFGKSGDDAKRFMRFGRDPTDKRFMRFGKSV*
>Cgig_Glycoprotein beta GPB5/GPA2
MWKRCVWLVVLAAFSAVCAVDLDSVTACLVREYNLFAQKPHVTPTGDVLECSGFV
KVNSCWGRCDSSEIADYKIPFKISNHPVCTYSRVQKRRVRLPNCHPEHPDPYYVVY
DALACSCRYCNSKYTSCETLNG
>Cgig_GNRH/AKH
MYTHRLTVLLLLICLTLGHAQWAQTFGWGGAGNGKRSAWSTSSNSEKYDCSQNN
ENILNVISSLVQLEVQRLNYCENKRRILTQ*
>Cgig_GNQQNxP-like peptide
MKLFVALLPFLGLVYCSAAPTDIEKRSAMVKRAQEVMMFGNQQNKPRIKKSDPEVP
ALPDLRGSVTANKAEDIAETKTAELPKAIEKELEDVSDDVMEPKESEDITSESGEPTT
DELLEDFPPETVEALKELENAKNDKEEKSSEEINNNASEKSSEENKEGEENSVQRT
DNDEDMLEVLPEEWYNNPALALQLYRAYQKLPNYDLPYYGRRRRSPLKARMFTND
MKRSHRNKRDLPYSDEEYPMAYYPSEFGPSFTLKDLEAFAREKEYEDSVLRTILDN
VGPDDVQEIEHQGVRGLFIPLEQEQVPVAPPSKRSSYFYPYSEEPETHFGAFVPEK
KEYLDTYSRLVQLARELSKSDSDKEDYQRWQ*
>Cgig_insulin-related peptide 1
MKWLLSVTLVSCLRVGVGSQLQACGSALTDILSLVCRNQFHAPAKRIDAPMVNDID
ELARYKRSRQFAVAQKRGGVVEECCFSSCSYENLLLYCSQSVDPLDVFTAARVGS
SSTTTQRPITTTSTPTATTTPLSVLTVTGHKPEDQAELDEKAAAAAAWALSRRWFY
VNRLGRFRGQVGRRIINM*
>Cgig_insulin-related peptide 2
MPVNRKHQIFILLCLHFTSVQSDFERVCNSQTDLRGPDPQGICGRLIPEMLHLVCG
GQYYVPSKRDVSSLSHHKQDRNVDFPRYSPLEGLILGKREASMYLTSQHSRTKRN
AYQGIVCECCYHGCNWFELQQYCGFRKKRNTEPDSISASSQNSGKLIDSVLNK*
>Cgig_insulin-related peptide 3
MTSETTWFIISLCLLQMVCPVLSGFEKVCTFETYRRGVHQQGACGDNLADMLRLVC
RKYKRSGGTRPGKTYDVLKRANFTAGGLSPPQIQARSAPWKSVLKRMMVSKEKAL
AFIANNLDFHRRKKKGSPYNLDKRYGDINIVCECCYHSCSVAEFEDYCAE*
>Cgig_LASGLVamide
MATTKSNAVLFTTIIYVVYTIGPIASQKSFRYDENLYQDQDETLVEKRQFDRLASGLI
GKRRLDSVASGLVGKRRLDTIASGLVGKRRLDSIASGLVGKRRLDSIASGLVGKRRL
DSIASGLVGKRRLDSIASGLVGKRRLDSIADGLVGKRKMDYHSNTYPDFPAAEKRMI
DSLASGLVGKRMMDSLASGLVGKRTMDSLASGLVGKRTLDSLASVIIYVVYTIGPIA
SQKSFRYDENLYQDQDETLVEKRQFDRLASGLIGKRRLDSVASGLVGKRRLDTIAS
GLVGKRRLDSIASGLVGKRRLDSIASGLVGKRRLDSIASGLVGKRRLDSIASGLVGK
RRLDSIADGLVGKRKMDYHSNTYPDFPAAEKRMIDSLASGLVGKRMMDSLASGLV
GKRTMDSLASGLVGKRTLDSLASGLVGKRQSFPDFSKGGNEED*
>Cgig_Luqin
MGLSEVMKFGELVTVFSVLFLVTFSMGDGAPQWRPQGRFGKRLDQRFPSPWQQI
SGTNKDIDVYPVLEEQTDVQANNVDNIINVLKKVCVESNVPGLFKCYRRTDSGFRS
SSGQP*
>Cgig_Luqin2

MKFGELVTVFSVLFLVTFSMGDGAPQWRPQGRFGKRLDQRFPSPWQQISGTNKDI
DVYPVLEEQTDVQANNVDNIINVLKKVCVESNVPGLFKCYRRTDSGFRSSSGQP*
>Cgig_Myomodulin
MLRLGRGLQMLRLGKRGMPMLRLGRSNGLSETDEDFMYPDESELDEGRRQVPLP
RYGKDLQQQLQLEWLQSVLDSDLENNGVRIIRPAGRPGRFRRSLKEDNGEEKDSE
ERHIPHPRIGRLIQLDDLKYPTSNLGSYYLTDSNVYTDKRGMPMLRLGRGMPMLRL
GKRLQADDSKRGMPMLRLGKRTNVQADSASQQETNQNKRGMPMLRLGRNAN*
>Cgig_Neurokinin1_NKY1
MPRYNNCSYVLCAITTIIFAIIATSESRDLSESDLMNFISTDQDRDFKVLSQLLKAEVL
KRKLRDLSLDSNDIDLSDELMNELSSEKRRRQYLPSLLTVRKRKVFWQPMPYVPHN
ARHNSRDKDAGASNDIPNSAAILRYG*
>Cgig_Neurokinin2_NKY2
MYRLNFHQIRTLAPALFLVLTFCSCAVSASRDDYQTEPTEREINGELAYILSNIRQKL
RSTDSDSYPVLLQPSSINAKRSRFGNPSTKKRGGIWIWMPAQGYVSVPRDEVGGA
SNKGSSSNLLRYG*
>Cgig_Neurokinin3_NKY3
MNISSAAMLRVFVVALWAIAILCQNTNGLTALENILAEETKDDYLRDVDDDSVLDQR
TREEDLKKLILKKLRFRELPNDSSSEVLNSLVKRVPDSFRYGDSLVDKVAALLRIKM
RPTKSPQVRMPSLRFG*
>Cgig_Neuropeptide Y/Neuropeptide F (NPY/NPF) 1
MQTSSLLAVLLVTLLSVWTVLGNDSLLPPNRPSRFSSPGQLRQYLKALNDYYAIVG
RPRFGKRDSEFSSYDQEPRSDGILSSSRDREDFAGPQWW
>Cgig_Neuropeptide Y/NeuropeptideF(NPY/NPF) 2
MHYKCNDAGRCDVILIDKGICYLLSSSIRTQQLIMKLFQSFIILVVLVDCVIVASHITKE
EDIQMLVPPVRPRIIATAEDLRRYLQQLNQFYMILSRPRYGRSVSKPVSEMAESS*
>Cgig_Neuropeptide Y/Neuropeptide F (NPY/NPF) 3 (PARTIAL)
MINCGFVTSHFTKEEDIQMLAPPERPRIIATSDDLRRYIQQLNQFYLILSRPRYGRSV
AKRIPKNHLQSISAMSN*
>Cgig_PDF/Cerebrin
MRTETCVTWVVCSLCLITLSVCRPIKDDEREAFASDNEERLLLAAKLLQQMQQRQY
GIRYYPRIPTHKRNLGTVDSLYNLPDLLYRGKR*
>Cgig_Pedal_peptide/Orcokinin
MFTHHKTLFSLMAVLLLSIAAAKSSPAPTDYNRSDTEDIETKEQRSGEETDESTELF
DDKQLGSMGTQEYSKKSMDSINGARGFRGFAKKHYDSIGGSRGLMGFNKKNYDSI
GGSRGLQGFNKRYYDSIGGSRGLQGFNKKNYDSIGGSRGLKGFNKRYDSIGGSR
GLKGFNKRYDSIGGSRGLQGFNKKNYDSIGGSRGFQGFNKKNYDSISGSRGLKGF
NKRYDSIGGSRGLKGFNKKNYDSIGGSRGLQGFNKKNYDSISGSRGFNKKNYDSIG
GSRGLKGFNKRYDSIGGSRGLKGFNKRYDSIGGSRGLQGFNKKNYDSIGGSRGFQ
GFNKKNYDSISGSRGLKGFNKRYDSIGGSRGLKGFNKKNYDSIGGSRGLQGFNKK
NYDSISGSRGLQGFNKKNYDSIGGSRGLQGFNKKSYDSISGSRGLQGFTKKGYDSI
SGSRGMKGFNKRHYDSINGSKGLSGFVKRSYDPISGSHGFSGFVKRETNTNGSKN
SESF*
>Cgig_Pedal peptide/Orcokinin 2
MWKLSFGAIKMQSLMNVLLVTLTISCCINRATGTADEERSSRHKRMLDRLGSGFIKR
EPDSKIDDLVPKLRYYLAKAISEQDDDSSPTEVDKRRLDRLDMGFIKKRRLDRLSTG
FIKRDDDTEYENENEDGEDQMDKRYLDRLDSGFLKRFVPGEDKRYFDRLGSGFIR
RSNFDRIGSGFIKRRFDRLGSGFIKRRFDRLNSGLIKRNLDRLNSGFIKRQRLDRLG
SGFIKREEMDKRRLDRLGSGFL
>Cgig_Pleurin

MQSLFWALIVLVSCDCALCLFYTSSKENDYPRFGKRNSRIHSDAWMDKDSIDDYNE
IIDSSEIKPTQYLSQSRLRMFKRGTNLLFQFLDRNGDQLITKNEFTANSFTEILDALTK
DHHGK*
>Cgig_Proenkephalin like
MLMLVFLGMFCLSAADKENPIPVSDTGEGTRCDADLNIAQTCIACSRLPQIIAVKLTD
CCSEDKAFQFCQICTKNSDDCLKEAFSINNLEKGVVLDGYSDVEDVPPEVIADNDM
NKRFGTLSMGGSSGYLYGKRDLDKRYGRLGFGGSRGFIYGKRSDDKRFSRLGISG
SSGYFFGKRDMGVDKRYKSLGMGGSTGFFYGKRSDDEKRFGTLGLGGSSGYLYG
KRDSDKEKRFWRLGIKGSNGYLYGGKRKRFGQLGMDSSSGYFYGKR*
>Cgig_Pyrokinin peptide partial sequence
MGRSAFYFPRMGRNQESADQEQSEEDIPCCNVGVSRAWGPEDPEPRRICPAAKC
CGHLSSKVYVDSDLTVYEICVSDVAALPMEQKS
>Cgig_RGWamide/APGWamide
MESPSLYLIVIALVLGIVSTDEELIDKRRPGWGKRGSLDFGSNNLPNSIIEKRKPGWG
KRGMEDEFEMNKRKPGWGKRAPGWGKRSFNEDLNNFIEKRRPGWGKRTSDLMD
ALNAMEVNKRKPGWGKRSEMEKRKPGWGKRTVPELSETDLSYSSIPTDSFSVLDK
RRPGWGKRSDSDISVERRAPGWGKRAPGWGKRSESSTCPDLEYQIQQLMNTILK
LEEYNRSHCGRQSPTSSNFGNSECEVVKIT*
>Cgig_SIFamide/NPFFamide
MKIYSIISIVIALVAVIVLKTSASKENSRGLTRLVGQQPLLFGRRGMNPNMNSLFFGK
RAVDRPTLDDIIVEKCSRIMAACQEYAHERMGEDDI*
>Cgig_sNPF_precursor
METKTLLIWVFAAFLAFIEVSTASKESKLNHEESEVKSLKRRSNPMDDVTDTYDFSD
EDLNELMKRGSLFRFGKRGALFRFGKRSEEDKRGSLFRFGKRMDGYYVSPYASSE
NFPEKDDKRGSLFRFGKRSSLFRFGRSVDNEKPHTPFRFGREEEDEI*
>Cgig_Sulfakinin/CCK
MKNSLEHLLISSVLVLCLLFTLTKGNSHAQNSIIQLSKLFSNLQDIHDAKQTQEENTK
DQVRSDTRKQRGEIGLILALGAPVHVESLQHNDKDTQTDTDKSAWDVDFEDDQQD
VEKRQGAWDYDYGLGGGRFGKRFDYNFGGGRWGRDVDHVKQGQAK*
>Cgig_Tachykinin
MVYLQRALFSLVFLITYSLTSATTGQLWDSVESDLDDTIESGDPVHKPLINFLINPKD
LHKAGIPESSSEEKRFGFAPMRGKRFNNDVEFPLVRNNKRARFFGLRGKRVPSLE
QRLSDEDFKNILGKLLNDAYDFSTNSNDADSSLEKRFRFTALRG*
>Cgig_TRH 1
MDAKFEITLYRENPIEMICSQFSGKGALLSSKEQTKAAGKADDRVKRSPFVPKFIGK
RDDENEAALLSLEAAIRDELLSQEPFDIYPSDADDAFEHEARMNRQLLVDKHDRNP
LGKRTAPILGFKRFSPREELKRSTILVPEDISEFDMNKRGNLPMFVGKRRAPLFVGK
RRMHLVVGRGGKMNNPKLVGKRGTPLFVGRRRADTDDKTIYYSYITLRRSVPDEL
RMRDSIADSLLNGDNFYSAADTGSGVLDIPDDEDSLSNSSSQEFKRFETPDCRSVT
IKRPVAAEFLSRSPNDSDQENLLSSAIRGLTETRIGTPDDPLLADSPLNIKGGYLENA
ETFDTDHLKLQNTLDENCSDSIHSPVIKDLSEELDNSDNQKMAENPTENQIFKKYSE
FLGKRYSEFLGKRGLPKRFSEFLGKRLGQIKRYSEFLGKRLNYKRYSEFQGKRNHV
QKRYSEFLGKRFYPRKRYSDFLMGKRYSDFLMGKRNIWSKKRQGEWVGK*
>Cgig_TRH 2
MKLTFGTILTLLWAFSSIKAKPAEEQTKAAGKADDRVKRSPFVPKFIGKRDDENDDA
LLSLEAAIRDELLSQEPFDIYPSDADDAFEREARMNRQLFVGKRDSNPLGKRTAPIL
GFKRFSPREELKRSTILVPEDISEFDMNKRGNLPMFVGKRRAPFFVGKRRMHLVVG
RGGNMNNPKLVGKRGTPLFVGRRRADTDDKPIYYSYIALRRSVPDELRMRDSIADS
LLNGDNFYSAADTGSGVLDIPDDEDSLSNSSSQKFKRFETPVFIGKRNSLIADLASA
KQAHSQLLQKRFEPPMYVDDSDQENLLSSAIRGLTETRLGTPDDPLLVDSPLNIKG

GDLENAETFDTDHLKLQNTLDENCSDSIHSPVIKDLSEELDNSDNQKMAENPTENQ
RSKKYSEFLGKRYSEFLGKRGLPKRFSEFLGKRLNDKRYSEFLGKRNHVQKRYSE
YLGKRFYPRKRYSDFLMGKRYSDFLMGKRNIWSKKRQWEWVGK*
>Cgig_Vassopressin/oxytocin
MECCKILHLSLPQVSLLLLMFTTLSSCCFIRNCPPGGKRSMGIVSQPTKECMSCGP
GLLGQCVGPDICCGPFGCQMGTSESNICGKENESTTACAISGPPCGSRNQGNCVA
DGICCDTGACSFNTKCKLNSEPRDVQILSLLKTLGLIETKAYNNAGMS*
>Cgig_Whitnin
MEISMPMLLCSSLVLVLISSVCSVPIDVARETENDHHLEEKRPKYMDTRDLYAFEKL
VYLALKGLVEEGKVNPEVMTSEANEGFLDNQGENGESKEKAGPADKRGHLRICVR
RSGSRYVPYPCFRS*
>Cgig_XHFAamide g10225.t1
MNRPMLQSLCVLAFCSTVLGAFSVQKFCENACNRGLGGNLCRCNGFHFAGKRTV
LENILPVPNTYSSVNGDEPINQRIQAPSFTNDIQQPAPVLRRENLSRIDKIRDTQMIIQ
WLLNNIRLIKSEMPEKHMDAYGIEDDQLTP*

**Table S3. Sequences of *Prosthecereaus crozieri* neuropeptide precursors.** The predicted signal peptide is shown in blue, predicted monobasic/dibasic cleavage sites are shown in green and predicted mature neuropeptides are shown in red.

>Pcro_7B2 peptide
MKNFLLALTICFVASNVGATNYDSFISELYDRNQRLMNDISSNQLPEALRRSMESEI
NDNHAFPIGPDAEDEEDEESDWGDRDDLIQKEEEIRNAEIPSQNNLWGEHRVSGG
TGKGSQSYLGDGITSKPEVKTNNLPAYCDPPNPCPIGMNPESATSPCDQHVEDTK
RFNQEWIKNKMKNGECPCDTEHMQHCESSYNRMKTGGRSNNPYFSGNKRAALA
AKKSPTMWTDERKNVETLQHSNPFLQGQAHAVKKKNGRFDRPYM*
>Pcro_Allatostatin-A 1
MNNRRRVYVWTSHCCLFLFVVIFKSSLGQLPLQDEFISPSVESAESSSNLKSENVP
VFLKDEVLERINLPEHLSKPVNSKSDFDTVTEENDPETTDLQKRDYLYSQYHKKAAL
RRRASTKYRRQAWYSGYGKRSFGQAWHSGYGKRGSSQAWFSGYGKRRRGQA
WFSGYGKRRRGQAWFSGYGKRAKAGQAWFSGYGKRGKAGQAWFSGYGKKRG
KAGQAWFSGYGKRRRGQAWFSGYGKRRRAGQAWFSGYGKRAKAGQAWFSGY
GKRGKAGQAWFSGYGKRGRPGQAWFSGYGKRGRAGQAWFSGYGKRDEDGQA
SVKGSWEARRK*
>Pcro_Allatostatin-A 2
MQSLTLNIIIFKLIALVVLSFEITSQSSFESTGMLSEDLDDNDLLQSPIHVVARRRAGS
FRSRLGYSSGYGKRASIRQRKESFRNRLGFSSGYGKRSMPSFWRQRRRLSFHNR
LGFSSGYGKRRMETRQRRKGSFRNRLGYSSGYGKRSDDLNS*
>Pcro_MIP/Allatostatin-B Wamide
MPKIFQLLLLCVLSIVAYASATDSVNEDHVISSRAAMGRLPWGKRAVMGRLPWGKR
AVMGRLPWGKRAVIGRLPWGKKAVMGRLPWGKRAVMGRLPWGKRNSDAEETEK
RGVWSNDAWGKRGVWSNDPWGKRGVWSNDPWGKRSDFDSEIDSDDDQKKRS *
>Pcro_Allatotropin
MSSSNRSTILSLTTSQYAILISCIFLLLLSIVPEISSASSLDGMEMKKSAGMNRRGSFK
NRLSFGDGFGKRSMPWADRYALRQRQSRVNHLASLMQELLDTMEE*
>Pcro_Bursicon-like
MQILSIFLMTLMPIINSQTGTKNNGEASTVTAMATNSIADILPGQRNTERAAHHLRHQ
RRHINSRPRLLDNLISRGGRRNRKRKTNTLRGFQMNRLQSLRRVWCQRGMFRQKI
KHSNCKKRTITNYLCYGQCSSVYIPQYRNRLTQRPQAAFQSCGFCFPKRTRVVRV
RLQCPRRNPPYKDKFVTAIDECRCIDIKSLRVHL*

>Pcro_Calcitonin
MSSKSVIFLIAIICIGLHSVVDGRTTQRRRRSTEEAELERFLPILDRMMAEQVKSQDP
ADIHTVLKRARCHFLNLGGLCNSAKAAEMVDQYRWLTRPGGPGK*
>Pcro_Corazonin
MIRCLTLALFTLVLVLVLDQTVGQNIHFSHDDGAGSSSWQAGKRAHQGHSLFRIKR
VYHFFRLRKNEQCVFPLALRKHIEQQMAEKTACSDDVDFASNLLKELEVKNEGR*
>Pcro_FVRIamide
MKPFIVLVFNGLPVALMVVLLSALASAADPPAVEEILQRQCIQNCIESYDLSIRSCQE
ACMQGDSGSRSADKRASFLRLGKREFLRLGKRASFLRLGRNFPSKKAVDQLDNA
MQKKASFLRLGRSGKEKKASFLRLGRESPDFKKVYKKATFLRLGRSSNNMLDDSEI
LN*
>Pcro_FMRFamide (no signal peptide, potentially an assembly error)
MVNVSDALSIFKSGKRSKEIPMADGLTRSPNEVGITRDKDFEYIALIDNTPLEMDEIR
NHSKASEEIMNIMEHISTGKKIKRGLYVQVCDRLSISHDVLFYGSQPVIPETLRKRAL
QLLHNGHPGMVQMKSYAKNTIYWPDINKECERCVLARDICQSKKTSAKPLQFESW
PKTNSFGDRIHIDYLGPIDAYMILIMGDSATGWINCDFTKSTTSDETIKMVRTFFAST
WSNGIAERAVRSFKERERTLKYINNIERRIDNLLLALNSGSNADMKLFGRKLWTPIT
NSHSTSDSWNQPTQLVWYRSFTNDNDIWMKGLADSVDSPSIRGVDEGGKEWRRS
IYHVKIRADDNEQHDEPSTTPRTDTEVEPEISSPNVDATFLVSISFSHHLTGAVDADT
DASDKKRASYIRFGKRSSGVPEIDDDKLIGNSDKRARYIRFGKRAGYIRFGKRSDDL
NGQIDLEDEGSNDKRARYIRFGKRARYIRFGKRADALYEGEHPDFGSDSKRASYIR
FGKRGRYIRFGKRARYIRFGKRDNEMNKEVDEKRARYIRFGRR*
>Pcro_GGRWamide
MALAFPFKRLEVSPIHPINQLDNLFNISQHFQLSQSPQSTLRPPVPALTFDEDTDKQ
ENPDYRRSVISILIKEGGKSPRAIVGNSMRRLMDEVILAQYNLEGKNKRKFAEIFCRL
IYDVVLSKEKENEKDGQRWGMSTYKEVLPMIRSVLKIANDKNARKGENVNGQKL
>Pcro_GNRH/AKH
MTSSYCVCFFTLFVTIYCSYTSAQTFHYSNAWRPGKRSNDMPRLSASLKDQLSSFE
HKECVLSHSTRAALWDQMFKNEPCVSDLNFLAAILDKRLRVISRK*
>Pcro_GNQQNxP- peptide precursor
MHLNLINPTHQYCLPHSPLRSYDTLALCFPFLFTIILRHDHPHPPNVKRADETVMFG
NQQNSHKTVKKSGPVPLNSKLMEKQAKVADSHAAAAMEDDPVKELAAMMASKPA
TKTQESEPVKDLHTLISQGDSEPNAPLIEEEKKRSSDLKENKDINMGDKPKRPQPIN
EDSDRQDYDDERYMYLLSKFGPGPLMLENESGSRRRKRAATTKLTGTRLNRIRRD
MSKMASESHMSMNSVHRKKRDIEDMDPDQLARLEYMLSRSNDGYEIPEEVVDPSY
AEDEVPLEYDPLEVERRGMLDDYSRYAVAKRVPDIEEAPEEVEEEQETDPVSVYES
EYGPVTYGLVEHEGVPGLFKRKTAMTSAPD*
>Pcro_IPRFamide
MFSGHLCNPRLIQLWLIGCLFIVNAHAWSAQTLTCSLGLTLIPQTWTLSDISESSKDK
KLDESFSDSSNARLSDQPEPLTADDMTYLPDEWQHDSSEIEEMDKRGFIPRFGKR
GFIPRFGKRGFIPRFGKRGFIPRFGKRGFIPRFGKRGFIPRFGKRGFIPRFGKRGFIP
RFGKRGFIPRFGKRGFIPRFGKRED*
>Pcro_LASGLVamide
MINLGLKLCLFSTICFIILVNFGVTTAGQRNSDTFDSTKNLTFATKSDEKDKLYGSES
PMKPNNRIQAYKRRGLLGKRSHTLPQDVDKRGDGNLYYGLLGKRESEKTSATDIA
RYYSNKQPVKSNDLELGLLARQPYKLSSELYGNFRKFSEKRGGNLYHGRLGKQGR
ILYTVPLRKQNRNLYRGLLGKRGGNLRRGLLGKRNGNLYGGLLGKRGGNPYDGLL
GKRNGNLYGGLLGKRNGNLYGGLLGKRGGNPYDGLLGKRNGNLYGGLLGKRNG
NLYGGLLGKRGGNPYDGLLGKRNGNLYGGLLGKRNGNLYGGLLGKRGGNPYDGL

LGKRGGNPYDGLLGKRNGNLYGGLLGKRNGNLYGGLLGKRGGNPYDGLLGKRNG
NLYGELLGKRNGNLYGGLLGKRNGNLYGGLLGKRNGNP*
>Pcro_LSDWN peptide
MFNIFRTIASFAIASVLFSVACGNGEENQSPRYATEENQGRIGNQPFYIHSVDQYKF
LHHQPPSKWTQSQVHHWKYKEKPLIRKRRAMLSDWAKRASLSDWNKRASLSSW
NKRASLSDWNKRASLSSWNKRASLSDWNKRASLSDWNKRASLSDWNKRASLSD
WNKRASLSDWNKRASLSDWNKRASLSDWNKRASLSDWNKRASLSDWNKRASLS
DWNKRASLSSWNKRGGLSDWDKPSPSSDRNLHKELADVLNADSRQYFGIVVPEE
SVKDDSNTDDKSDQGLAS*
>Pcro_Myomodulin-like
MALHIFTLFSIILCALSITVASPSNSEEADSESNAEYIDYHPGHTLRDLLNMAEDDTYL
PEPLRRTYKRAVRLIRLGKRDAEKRAVRLIRLGKRDMDKRAVRLIRLGKRDMDKRA
VRLIRLGKRDMDKRAVRLIRLGKRDMDKRAVRLIRLGKRDMDKRAVRLIRLGKRDM
EKRAVRLIRLGKRDMDKRAVRLIRLGKRDGDQQCLLQGNHDEVKQLANMMHEDD
KRDVRLIRLGKRDSDDDDEYCVTVEKRLVEDSSSKDEFGDKKGKPIIRVGRVG*
>Pcro_Myomodulin-like2
MTPLSFIVTLGLALICACAADENRVKRQVVLPRYGKEANYDLMEELKDDIKDDSDRS
VMLPRYGKDEEPEQFEEKRAIRLMRLGKRDSDEKRAIRLMRLGKRFSTADKRAIRL
MRLGKRYNIDEKRAIRLMRLGKRFDEKRAIRLMRLGKRNSKRAIRLMRLGKRSAES
DE*
>Pcro_ Neurokinin_NKY1_consistent
MVTRYHAAFLLFTIIVTVVTSSPLVRMKRSKAYERELYKGLERNGDNEFDRIRRIDID
AGFRSRFDAGSSFGHAKHAADLVRMYG*
>Pcro_Neuropeptide Y/Neuropeptide F (NPY/NPF)
MTRRMAFLQSLIVWSLAILLLVCATANASEPVERRPSGSRKFKYFQNSDQVKRYLL
TRLVAYYMQYGRPRFGKRSNPSSEYYPPERDTTYVTEYN*
>Pcro__Neuropeptide Y/Neuropeptide F (NPY/NPF) 2
MCLRCLPVLFNSSRSRISWSKMLIQLFAALFIFICIDDHLTEAAVVQRRRPRFFSNVR
ELQAFIAKLNLYYQIQGRPRFGKRSMLQDRFFT*
>Pcro_Neuropeptide Y/Neuropeptide F (NPY/NPF) 3
MPLLRSCSRFLPLGIIIQFILVLRAQDSCGMRFRPPPPPADVTSKDQVAAYLSALGN
YFRMVERPRFGKRNFVRRSDIETESTF*
>Pcro_RGWamide/APGWamide
MSYSLTFYWNLTVFVLGLTTILIATASGDDGLGELDPGIEYAELIFEDGDYGGYANKR
SPDKRAPGWGKRAPGWGKRSPDKRAPGWGKRSPDKRSPGWGKRSPDKRAPG
WGKRSPGWGKRSPGWGKRSPDKRAPGWGKRSADRRSPGWGKRW*
>Pcro_RGWamide/APGWamide2
MARTKQTARKSTGGKAPRKQLATKAARKSAPATGGVKKPHRYRPGTVALREIRRY
QKSTELLIRKLPFQRLVREIAQDFKTDLRFQSSAVSALQEESEAYLVGLFEDTNLCAI
PAKRLTIMPKDIQLARCIRASGDDGLGELDPGIEYAELIFEDGDYGGYANKRSPDKR
APGWGKRAPGWGKRSPDKRAPGWGKRSPDKRAPGCGKRSPDKRAPGWGKRSP
DKRAPGWGKRSPDKRAPGWGKRSADRRSPGWGKRAPGWGKRR*
>Pcro_sNPF
MILPTERDILLNILDAVVSLISHCKLHHDWNLGGDIAVVSRYLLQSGLEWTTITLPGCS
SAVVAVDYQGVLVFYIVPVLIFTELLSTFRKTPAYTADFVRMLKETLSVVIAFIAMCAL
TRTDAVSSKTKRDDYPSNLFRWGKRSARSEILSEVLRNYVDEFGDEGLKKRNNLF
RFGKRRVHTPFRFGRELDEE*
>Pcro_Sulfakinin/CCK
MHLSSSSFLYLPQRLCNLLLLSFLVLVLHSCQQTCHTAPAKSHARSMALLAPLLRQR
FALDDMIHAEKRMRDEKKDSEDYFEGSDYGWAGGRFGRR*

>Pcro_TRH1
MMHKIYLIFASTLYLTIGIHGSSSRGYVSNQPVDDIEKRVSEFIGKRSLLSQLIEDYRK
FGAKSPKKRYGEFLGKRTGEFLGKRDQSSLISSMNSRMAPDRSVYYSGGEFLGKR
KKKFIAKRFGEFLGKRATPYSLRNKQTFPENGLDADAFHNKRVSWPN*
>Pcro_TRH2
MRMSNKARCFTLAVVISIAFYLSIQFRDETKINNDVKKWIYTLIYMDKSRFQRIETKPS
NQRIESSIANSAKIEISTTVKGVAYPSSNGTPVVSTNHRKINTIVEENAKHKHNSIKKK
KKNVQKPPSMYAKMRRQVACRRTTEKFFLNAKKTKYGIPHIIHQIYKSAIIPVQYVHL
LDKCRRINFDFIHVIWTDKKSEELLYHHYPEYKSLLQQYKGIQKADFFRILILYHYGGI
YMDMDVECRKPLENYINGLHSGAVFDQERRLQTRLLWNLDYLTMNSVIITPRKHPF
FKYLLDHFKDTLEIKNVLWKTGPFYVHGRLMEYQRQNKTGSHRVSLADPKDFSPLL
DRTHFKNFNAKCSGNITALSIFQQRGCIEWLKQGRKQGDIRQLTKDSIGVHRFLHLD
QPVDDIEKRVSEFIGKRSLLSQLIEDYRKFGAKSPKKRYGEFLGKRTGEFLGKRDQS
SLISSMNSRMAPDRSVYYSGGEFLGKRKKKFIAKRFGEFLGKRATPYSLRNKQTFP
ENGLDADAFHNKRVSCLIRIDANLKFPSSDEITSNFCKVL*
>Pcro_Vasopressin/Oxytocin
MKSVMASSSRPRRLCAEQALRGILSGVDEDDVNEAADDATESESDHILTEDEESSD
NEDIDIPDELPSDELPSTVPVLLLEEMGWSGLQHVQWYDEGELLTLTMSMMPIIELL
SLMLIVSSCSGCFIRNCPPGGKKRSTGIPESSHTIRECTSCANGRGHCVGPDICCG
KTIGCQMSNQISSVCQRENEFPTPCKSKGKRCGLYNIGKCSTSGLCCTSEELSDIYT
SPAEVGPPLSNSDHNSVLLTPSSQPIIDNTVTKTVFDLRRSNIDHFVSRLASSGFDK
VYAASDVDDKCHAFYELFFDALSVIPHHEVIFTKKDKPWLTPILKKMINDRWRAYRV
QNWTVFNHLKTKIRTEIFKAKKTWSERVMRKKKNVWNVVKEFQGSNKASTSFPDS
NIELSRFLSDLSDTFASCFNNEKDEDLCGLSDEIWDLHFDSYDVYNSLKRLKFRQST
GYDGISARILWECADIICEPLFEIYRTSVSTRVFPEYWKNAFIVPTPKRSSPTIHDYRQ
ISLLPIIANVFERLVLASMKGLFLEFFGSSQHAYRPYGSTTSALIEIHNYITLFMECPN
NLTVKITCLDYSKAFDKLQFNRLLNNLLRWNFNRGFIVWLSSYLRNRCERVRIAREI
GPVIDIPSGVPQGSVLGPYLFALFIATLNIDSPVAKLVKYADDLTLIESCSIDNPCPNN
LEHVLMWSCQNKMPLNQKKCQQILIHRVRNDSAVPYDKN*
>Pcro_XHFamide
MNIFSCCIFVLSGLGILIQRCNCDAARRESIGSLGDNLIRFGRFTKGVKGQANDEHQ
KRRTSKRYFGDNILHFGKRNVKRYYGDDMLHFGKRDSKRYYADDMIHFGKRNAKR
YYGDDMLHFGKRDSKRYYADDMIHFGKRNAKRYYGDDMLHFGKRDSKRYYADDM
IHFGKRNAKRYYGDDMLHFGKRDSKRYYADDMIHFGKRNAKRYYGDDMLHFGKR
DHSKRYHPDDKLHYGKRSAKRYYGDDMLHFGKRDHSKRYHPDDMLHYGKRNAK
RYYGDDMLHFGKRGNLKRYYGDDMLHFGKRDSKRYYGDDMLHFGKRDSKRYYG
DDILHFGKRDSKRYYGDDMLHFGKKEKSSEIDDDTSN*
